# A conserved *HOTAIRM1-HOXA1* regulatory axis coordinates early neuronal differentiation

**DOI:** 10.1101/2022.08.17.504180

**Authors:** Dana Segal, Samy Coulombe, Jasper Sim, Josée Dostie

## Abstract

HOTAIRM1 is unlike most long non-coding RNAs in that its sequence is highly conserved across mammals. Such evolutionary conservation points to it having a role in key cellular processes. We previously reported that HOTAIRM1 is required to curb premature activation of downstream *HOXA* genes in a cell model recapitulating their sequential induction during development. We found that it regulates 3’ *HOXA* gene expression by a mechanism involving epigenetic and three-dimensional chromatin changes. Here we show that HOTAIRM1 is required for proper progression through the early stages of neuronal differentiation. We found that it associates with the HOXA1 transcription factor and participates in its downstream transcriptional program. Particularly, HOTAIRM1 affects the *NANOG*/*POU5F1*/*SOX2* core pluripotency network maintaining an undifferentiated cell state. HOXA1 depletion similarly perturbed expression of these pluripotent factors, suggesting that HOTAIRM1 is a modulator of this transcription factor pathway. Also, given that binding of HOTAIRM1 to HOXA1 was observed in different cell types and species, our results point to this ribonucleoprotein complex as an integral part of a conserved *HOTAIRM1-HOXA1* regulatory axis controlling the transition from a pluripotent to a differentiated neuronal state.

## Introduction

Transcription of long non-coding RNAs (lncRNAs) inherently points to potential biological significance; yet predicting their roles solely based on sequence has been challenging because of the rapid evolutionary emergence and decay of their genes, their poor primary sequence conservation, and lack of detectable conserved secondary structures [1-3]. However, the *HOTAIRM1* lncRNA gene is exceptionally conserved across mammals, with orthologues identified simply by comparative genomic sequence analysis [4]. *HOTAIRM1* has also been found in other vertebrates, including birds and reptiles, using combinations of short sequences and secondary structures [5]. Through microsynteny analyses, Garcia-Fernandez and colleagues even identified amphioxus *HOTAIRM1*, indicating that its origin pre-dates the appearance of extant chordate lineages more than 500 Ma ago [6].

*HOTAIRM1*’s syntenic conservation across distantly related species suggests that its position and co-localization with *HOXA* genes is a functionally relevant ancestral feature. Further bolstering this hypothesis is our finding that human *HOTAIRM1*, encoded between *HOXA1* and *HOXA2* (Figure 1A), localizes at the anchor point of a chromatin loop with the *HOXA4/5/6* genes, and that its expression is required to break this looping contact so as to curb premature activation of verging *HOXA* genes during neuronal differentiation [7]. Evolutionary conservation of HOTAIRM1’s function is further supported by finding that its disruption during *Xenopus tropicalis* development results in a severe headless phenotype, which is proposed to occur at least partly through interference of *HOX* gene regulation, and by extension, the transcriptional programs in which they take part [6].

**Figure 1.**
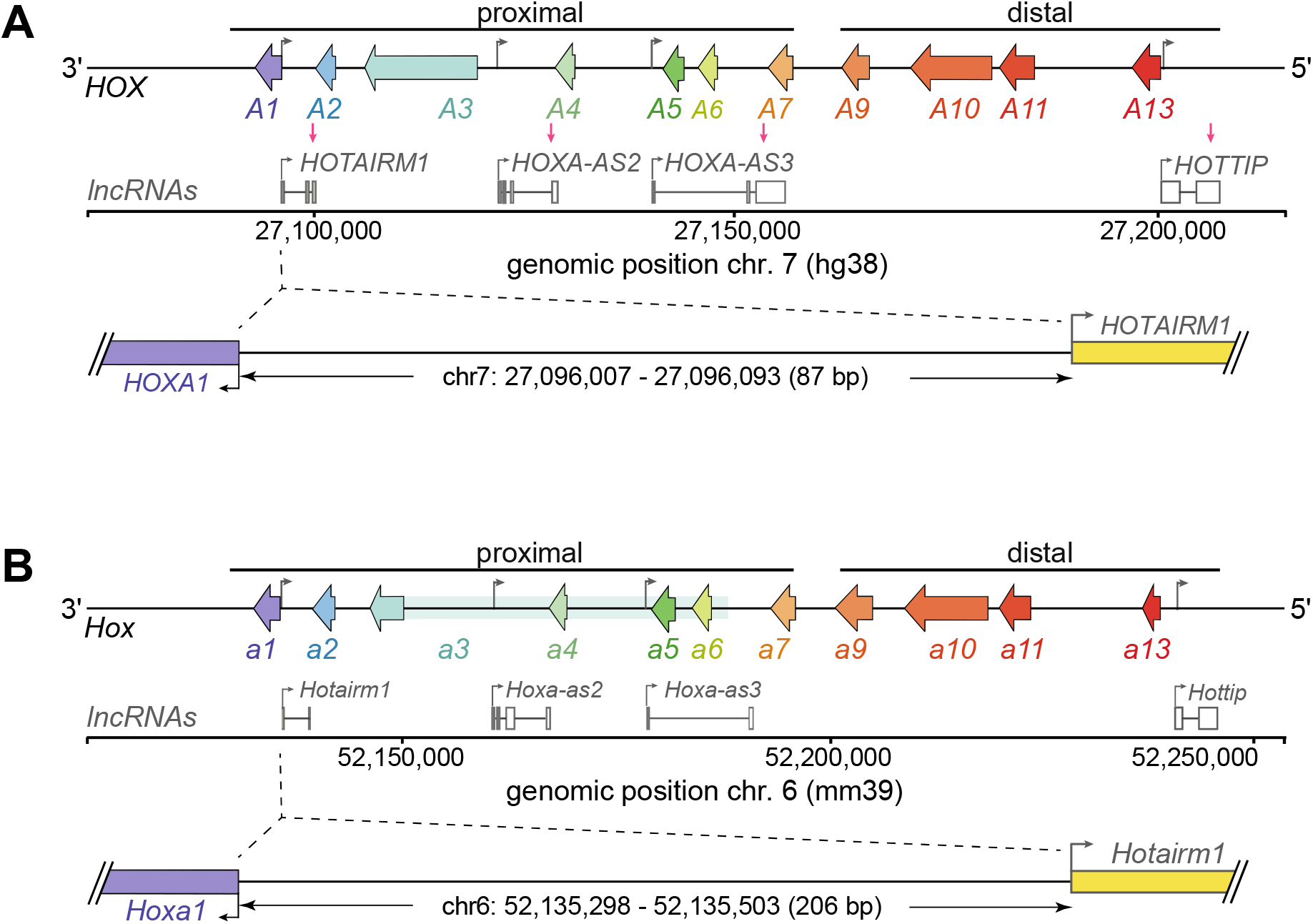
Schematic of the human (**A**) and mouse (**B**) *HOXA* gene clusters. Left-facing arrows represent *HOXA* genes and lncRNA genes are below each cluster. The shared *HOXA* and *HOTAIRM1* promoter region is expanded below the clusters.

The regulatory response of *HOTAIRM1* to retinoic acid (RA) is also conserved [7-9], as RA rapidly induced *HOTAIRM1* in human myeloid cell lineages where it was first identified [8], human pluripotent cell lines ([7,10] and results below), mouse embryonic stem cells (mESCs), and during mouse embryonic development [9]. Interestingly, HOTAIRM1 appears to feedback onto RA-induced gene expression programs of different cell types, as was reported in myeloid lineages where its RNAi knockdown altered granulocyte maturation and G1 growth arrest [11], and as we have shown during neuronal differentiation [7]. Furthermore, HOTAIRM1 was found to impinge on the regulation of *NEUROG2*, encoding a proneural transcription factor transiently expressed at the transition from neuronal precursors to differentiating neurons [10].

Given the rapid induction of HOTAIRM1 by RA in many cell models ([7,10,12], and results below), it stands to reason that it likely functions at earlier stages of neuronal differentiation as well. We thus wondered which key genes, beyond those at the *HOXA* cluster, HOTAIRM1 might influence during this fundamental process. Here, our findings reveal that a conserved *HOTAIRM1-HOXA1* regulatory axis contributes to mediating the transition from pluripotent to differentiated cell states in the early stages of neurogenesis.

## Materials and methods

### Cell culture

The NTERA2 clone D1 cell line (NT2-D1) is from the American Type Culture Collection (ATCC; CRL-1973). NT2-D1 are human pluripotent embryonal carcinoma cells derived from a lung metastasis in a 22 year-old Caucasian male with testicular cancer [13]. They were cultured in Dulbecco’s Modified Eagle Medium (Gibco™) supplemented with 10% Fetal Bovine Serum (FBS; Gibco™) and 1% penicillin/streptomycin (Gibco™). Gene induction and cell differentiation were performed by adding all-trans retinoic acid (RA; Sigma-Aldrich) to 10 μM. Cells were collected at various induction times as specified in the figures. The NCCIT cell line was kindly provided by Dr. J. Teodoro (McGill, ATCC; CRL-2073). They are human pluripotent carcinoma cells derived from a mediastinal mixed germ cell tumor in an adult Japanese male [14]. They were cultured in Roswell Park Memorial Institute (RPMI) 1640 medium (Gibco™) supplemented with 10% FBS. Gene induction and cell differentiation were performed by adding RA to 10 μM. Cells were collected at various times induction times as specified in the figures. The P19 cell line was a kind gift from Dr. M. Featherstone (Nanyang Technological University, ATCC; CRL-1825). P19 is a pluripotent mouse embryonal carcinoma line that can be differentiated into cell types of all three germ layers, including neuronal cells, by RA [15]. P19 were cultured in Minimum Essential Medium α (MEMα) (Gibco™) containing 2.5% FBS (Wisent) and 7.5% Bovine Calf Serum (BCS; Wisent). They were induced with 1 μM RA as specified in figures. All cells were grown at 37°C in 5% CO_2_ atmosphere.

### Plasmids and transient transfections

Flag-GFP (pcGFP-3XFlag-myc), Flag-hHOXA1 (pchHOXA1-3XFlag-myc) and Flag-mHoxA1 (pcmHoxa1-3XFlag-myc) were generated by cloning PCR-amplified inserts into a pcDNA3.1 vector (Addgene) featuring a 3XFlag-myc tag sequence in the multiple cloning site. Specifically, to build pcGFP-3XFlag-myc, forward (5’- GACGGATCCATGGTGAGCAAGGGCGAGGAGCTG-3’) and reverse (5’- CTGGAATTCCTACTTGTACAGCTCGTCC-3’) PCR primers containing BamHI and EcoRI restriction sites, respectively, were used to amplify eGFP from pPrime-HA-eGFP, which itself was constructed by subcloning an HA tag into pPrime-CMV-eGFP-mir30PGK-Puro, kindly provided by Dr. S. Huang (McGill). To construct pchHOXA1-3XFlag-myc, forward (5’-CGCGGATCCATGGACAATGCAAGAATGAAC-3’) and reverse (5’- CCGGAATTCGTGGGAGGTAGTCAGAGTG-3’) PCR primers featuring BamHI and EcoRI restriction sites, respectively, were used to amplify hHOXA1 from pcHA-HOXA1. The pcHA-hHOXA1 construct was generated by subcloning HOXA1 from a cDNA sample, reverse-transcribed using oligo-dT, from NT2-D1 cells treated with RA for 7 days. To construct pcmHoxa1-3XFlag-myc, PCR was used as described above to amplify mHoxa1 from the pBS31-mHoxa1-3FMS plasmid, kindly provided by Dr. R. Krumlauf (Stowers Institute; [16]).

For RIP experiments in NCCIT (6 million) or NT2-D1 (7 million) cells, constructs (12.5 μg) were transfected using Lipofectamine™ STEM transfection reagent as recommended by the manufacturer (Invitrogen™). RIP samples in HEK293T cells were generated from 40 million cells transfected with 25 μg of plasmids and Lipofectamine™ 2000 transfection reagent (Invitrogen™). P19 transfections were with 7 million cells, using 12.5 μg of plasmids and Lipofectamine™ 2000. ChIP samples in NCCIT were from 4 million cells transfected with Lipofectamine™ STEM and 12.5 μg of plasmid.

### RNA interference

RNA interference (RNAi) knockdowns were performed by reverse transfection using the Lipofectamine™ RNAiMAX transfection reagent and either 45 nM (NT2-D1) or 30 nM (NCCIT) of small interfering RNA (siRNA), as per the manufacturer’s recommendations (Invitrogen™). For transfections in 6-well plates, 600,000 cells (NT2-D1) or 500,000 cells (NCCIT) were seeded in each well while transfected with 7.5 μL of transfection reagent. Larger scale knockdown samples were produced by reverse transfecting 5 million cells with 35 μL of transfection reagent in 10 cm plates. The control siRNA (siNC; “*Silencer*® *Select Negative Control No. 1 siRNA*”) was purchased from Invitrogen™ (cat. no. 4390843). HOTAIRM1 siRNAs (siM1-1: 5’ - GAAAGGCGAGCTTGGTTACGCTTAA- 3’, siM1-2: 5’ -GACTTCGAAGCATTAACGATC- 3’, siM1-3: 5’ -CTAGAGTGACTGAACCAGATT- 3’, siM1-4: 5’ -ACTGGTAGCTTATTAAAGATT- 3’) and the HOXA1 siRNA (5’ - AGAACTTCAGTGCGCCTTATT- 3’) were purchased from Invitrogen™ (Silencer® Select siRNA; Thermo Scientific). RA was added to 10 μM 6h after transfection to induce neuronal differentiation, with cells collected at various times as specified in the figures.

### Live cell imaging and analysis

For the RA induction time-course (Figure 2B-E), single-cell suspensions were seeded into 6-well plates at a density of 200,000 cells per well. Cells were left to adhere for 6 hours before adding media containing RA for a final concentration of 10 μM. For RNAi knockdown samples (Figure 2G-I, Supplementary Figure S1), cells were reverse transfected as described below in 6-well plates at a density of 450,000 (Figure 2G-I, Supplementary Figure S1A-D) or 600,000 (Supplementary Figure S1E-G) cells per well. Cells were incubated for 3 days while imaged by an IncuCyte® Zoom Live-Cell Analysis System (Essen BioScience) every 4 hours, starting 30 min to 3 hours after adding RA as specified in figures. Cell confluency was approximated by the Incucyte® Zoom 2016B Software from 16 evenly distributed areas within each well. To account for irregular cell distribution due to variations in plate coating, the 4 central areas were omitted from estimates.

**Figure 2.**
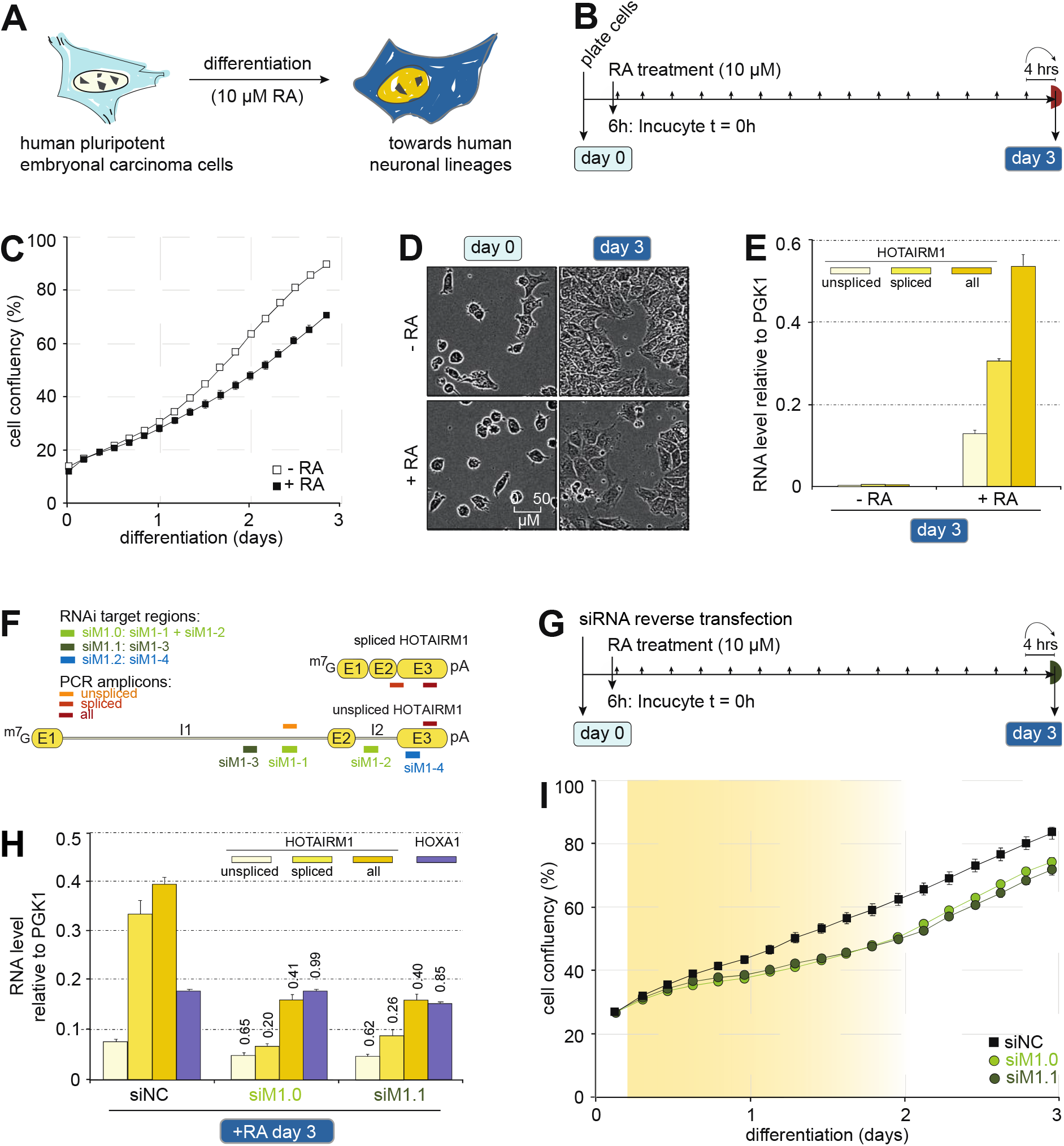
Depleting HOTAIRM1 hampers cell proliferation during neuronal differentiation. (**A**) Schematic of the method used to differentiate cells. (**B**) Outline of the live cell imaging procedure used to track cell confluency. (**C**) NT2-D1 confluency changes in the absence (-RA) or presence (+RA) of retinoic acid as captured by live cell imaging. Timepoints reflect when pictures were taken as indicated in (**B)**. (**D**) Representative phase contrast microscopy images of -RA or +RA NT2-D1 cells at the start (Day 0) and end (Day 3) of the differentiation time-course monitored in (**C**). (**E**) Steady-state levels of HOTAIRM1 lncRNA variants of collected samples monitored in (**C**). (**F**) Diagram indicating HOTAIRM1 regions targeted by siRNAs or quantified by RT-qPCR. 5’ to 3’ directionality is indicated by 7-methylguanosine cap (^m7^G) and poly(A) tail (pA); “E” and “I” represent exons and introns, respectively. (**G**) Outline of the live cell imaging procedure (*top*) used to track cell confluency (*bottom*). (**H**) Steady-state levels of HOTAIRM1 variants and HOXA1 in RNAi knockdown samples monitored in (**I**) and collected after 3 days of RA treatment. HOXA1 expression values were divided by 10 to display on the HOTAIRM1 scale. (**I**) Confluency changes of NT2-D1 cells transfected with siRNAs targeting HOTAIRM1 or a negative control. Timepoints reflect when pictures were taken post RA treatment as indicated in (**G**). Yellow shading highlights exponential growth delay when HOTAIRM1 is depleted. Error bars in cell confluency graphs (**C, I**) show the standard error of the mean (sem) across 12 culture dish regions, whereas error bars in expression level graphs (**E, H**) are the standard deviations (stdevs) between at least 3 RT-qPCR measurements. Numbers above histogram bars are fold differences relative to corresponding siNC levels.

### RNA extraction and quantitative real-time polymerase chain reaction (RT-qPCR)

TRIzol® reagent was used to extract total RNA from cells as recommended by the manufacturer (Invitrogen™). The resulting RNA pellets were resuspended in RNAse-free water and quantified using a NanoDrop™ 2000 Spectrophotometer (Thermo Scientific™) before 10 μg of total RNA was treated with DNaseI for 15 min at 37°C, as per the manufacturer (New England Biolabs Ltd.). DNaseI-treated total RNA was re-extracted with TRIzol®, resupended in RNAse-free water, and re-quantified by NanoDrop™. One microgram of each DNase-treated total RNA sample was reverse transcribed using SuperScript™ III and oligo(dT)_12-18_ (Invitrogen™) to measure steady-state gene expression with RT-qPCR as we previously described [17,18], using the ΔΔC_t_ method to normalize expression to Phosphoglycerate Kinase 1 (*PGK1*) or Actin (*ACTB*). The sequences of primers used for RT-qPCR are found in Supplementary Table S1.

### Western blotting

Whole cell extracts from RNAi knockdowns and lysates from RIP assays or were denatured 5 min at 95°C in SDS sample buffer (62.5 mM Tris-HCl pH 6.8, 2% SDS, 7.5% glycerol, 5% β-mercaptoethanol, 0.04% bromophenol blue). Protein samples from RIP (∼100,000 cells or 1%) or RNAi experiments (∼150,000 cells) were resolved by sodium dodecyl sulfate polyacrylamide gel electrophoresis (SDS-PAGE) before transferring onto 0.45 μm nitrocellulose membrane (Thermo Scientific™) with a Hoefer TE77X semi-dry transfer unit at 20V for 1 h. Membranes were blocked in 5% skim milk in PBST (PBS containing 0.1% Tween-20) for 2 h at room temperature. For RIP sample analysis, immunoblotting was conducted using the mouse anti-Flag antibody (Sigma-Aldrich, F1804) to probe transfected Flag-HOXA1 or Flag-GFP, and with antibodies against eukaryotic translation elongation factor 2 (eEF2) (New England Biolads Ltd., 2332S) as control. The RNAi knockdown samples were probed with antibodies against HOXA1 (Abcam, ab230513), NANOG (Abcam, ab109250), SOX2 (Abcam, ab92494), OCT4 (Abcam, ab181557), or eEF2 as loading control. Primary antibodies were diluted in 5% skim milk in PBST according to the manufacturer’s recommendation, and incubated overnight at 4°C. The membranes were washed 3 times 5 min in PBST before incubating with horseradish peroxidase-conjugated AffiniPure goat anti-rabbit IgG (cat. no. 111-035-003) or rabbit anti-mouse IgG (cat. no. 315-035-003) secondary antibodies (Jackson ImmunoResearch Laboratories Inc.) for 2 h at room temperature. Membranes were washed at room temperature 3 times 10 min in PBST before using chemiluminescence to reveal protein bands with the Western Lightning® Plus-ECL Enhanced Chemiluminescence reagent (PerkinElmer Inc.) as per the manufacturer’s instructions.

### RNA immunoprecipitation (RIP)

Cell pellets of transiently transfected, RA-induced NT2-D1, NCCIT, P19 or actively growing HEK293T were lysed in 1 mL lysis buffer (10 mM HEPES-KOH pH 7.5, 200 mM NaCl, 2.5 mM MgCl_2_, 0.5% NP-40, 0.5% Triton X-100, 0.5 mM DTT, 1X protease inhibitor cocktail (Sigma-Aldrich), 1 mM PMSF, and 0.1 U/mL RiboLock RNase Inhibitor (Thermo Scientific™) by incubating on ice 15 min, shearing through a 23 G needle (20 strokes), sonicating on a Covaris® M220 Focused-ultrasonicator (50W PIP; 10% Duty Factor; 200 Cycles per Burst; 7°C bath temperature; 30s time), and shearing again with a 23 G needle (20 strokes). Lysates were centrifuged at 14,000 rpm 15 min at 4°C to pellet debris.

Each RIP used the equivalent of 5 million cells in a final volume of 500 μL, with 10% set aside as input (for western blotting or RT-qPCR). Cell lysates were incubated with 5 μg of antibodies against the Flag epitope (Sigma-Aldrich, F1804) or a control IgG (Abcam, ab12073) overnight at 4°C on an end-over-end rotor. Magnetic Protein G beads (Dynabeads™, Themo Scientific™) were used to pull-down RNP complexes, as per the manufacturer’s instructions (, cat. no. 10003D), and were washed with 500 μL cold RIP buffer 3 times. RNA was extracted using TRIzol® as recommended by the manufacturer (Thermo Scientific™) and treated with 1 unit of DNaseI (New England Biolabs, Ltd.) for 15 min at 37°C. Samples were re-extracted with TRIzol®, resupended in RNAse-free water, and reverse transcribed with SuperScript III and oligo(dT)_12-18_ (Thermo Scientific™). RNA levels were measured by RT-qPCR using primers listed in Supplementary Table S1.

### Chromatin immunoprecipitation (ChIP)

ChIP experiments were conducted as described previously [17] with lysates from transiently transfected NCCIT induced for 3 days with RA, sonicated with a Covaris® M220 Focused-ultrasonicator (75W PIP; 20% Duty Factor; 200 Cycles per Burst; 7°C bath temperature; 10m time). Each ChIP was from 5 million cells, and 5 μg of either anti-Flag antibody (Sigma-Aldrich, F1804) or control IgG (Abcam, ab37415). Magnetic Protein G Dynabeads™ are used for pull-down according to manufacturer’s instructions (Thermo Scientific™). Purified genomic DNA (gDNA) from 10% of input chromatin is used for normalization. Final input and ChIP DNA were re-suspended in 50 μL of 1X Tris-EDTA pH 8, with nine Flag pull-down pellets pooled for RT-qPCR.

### Databases URLs

The ChIP-seq peak tracks for Flag-mHoxa1-expressing KH2 mESCs, without RA treatment or after 24h RA treatment, contain Hoxa1-bound regions consistent in occupancy between two independent experiments. They were published in [19], can be found at https://genome.cshlp.org/content/27/9/1501/suppl/DC1, and correspond to Tables S1 and S5, respectively. The NT2-D1 and H1-hESC ChIP-seq datasets for H3K4me1 (wgEncodeEH000917, wgEncodeEH000106) and H3K27me3 (wgEncodeEH000908, wgEncodeEH000074) were obtained from and visualized in human genome assembly hg19 using the UCSC genome browser (http://genome.ucsc.edu).

## Results

### Depleting HOTAIRM1 hampers cell proliferation during early neurogenesis

We started examining the role of HOTAIRM1 during neuronal differentiation in the NT2-D1 human embryonal carcinoma (EC) cell line [20]. These cells are distinct from most EC lines in that the only identifiable retinoic acid (RA)-differentiated phenotype is neuronal (Figure 2A) [13,21]. We first used live cell imaging to visualize the impact of RA on cell growth early during neurogenesis (Figure 2B). As expected when differentiating cells exit the cell cycle, RA treatment abated exponential growth, as measured by confluency (Figure 2C) and confirmed through visual inspection (Figure 2D). When determining HOTAIRM1 RNA levels, we considered its spliced and unspliced forms, as both were found in NT2-D1, with functional implications in *HOXA* cluster regulation [7]. As expected, both HOTAIRM1 forms were not significantly expressed before differentiation and were potently induced by RA (Figure 2E, Supplementary Table S1) [7,12].

We next imaged RA-induced cells transfected with siRNAs targeting HOTAIRM1 (siM1.0, siM1.1) compared to a non-coding negative control (siNC) (Figure 2F,G). Both siM1 targeting designs considerably reduced the overall levels of the transcript at Day 3 post RA (Figure 2H). We found that depleting HOTAIRM1 slowed the rate at which cells increase confluency after RA induction (Figure 2I). Onset of this delay occurred within hours, in line with HOTAIRM1’s rapid and potent induction by RA and an early role in differentiation ([12], and results below). Cell growth was hampered until at least Day 2, and this effect was reproduced across biological replicates (Supplementary Figure S1). As NT2-D1 do not grow evenly in monolayers and shift morphology during differentiation, automated cell feature analysis is unreliable. However, manual parsing of live cell images from differentiation time lapses revealed no considerable difference in cell death, cell size, or morphology, indicating decreased proliferation upon HOTAIRM1 depletion underlies the observed discrepancy in confluency (Supplementary Figure S1B). We verified lower cell density as proxy to cell count by overlaying a grid onto each image, and averaging cell numbers from 4 areas containing each at least 600 cells within the first 48 hours. We detected lower relative cell counts in HOTAIRM1 knockdown samples (Supplementary Figure S1C,D), an effect not sensitive to seeding density (Supplementary Figure S1E-G), altogether indicating that HOTAIRM1 is necessary for normal cell proliferation early in neurogenesis.

### HOTAIRM1 depletion interferes with proper neuronal differentiation

To understand why depleting HOTAIRM1 affects cell proliferation in early neurogenesis, we examined whether differentiation itself is impacted. The SOX3 (sex determining region Y-box protein 3) transcription factor is amongst the earliest neuronal differentiation markers in vertebrates [22]. It is a member of the SOX (SRY-related HMG box) family regulating stem cell identity: SOX3 is transiently expressed in neural progenitors to help maintain an undifferentiated cell state and counteract proneural proteins [23]. In NT2-D1, we observe transient induction of *SOX3* by RA peaking around Day 2 to 3 concurrent with *HOTAIRM1*’s expression pattern (Figure 3A). Upon HOTAIRM1 depletion, we consistently measured SOX3 expression at much lower levels relative to control samples (Figure 3B,C) in 3 biological replicates using two distinct siRNA targeting designs (Figure 2F).

**Figure 3.**
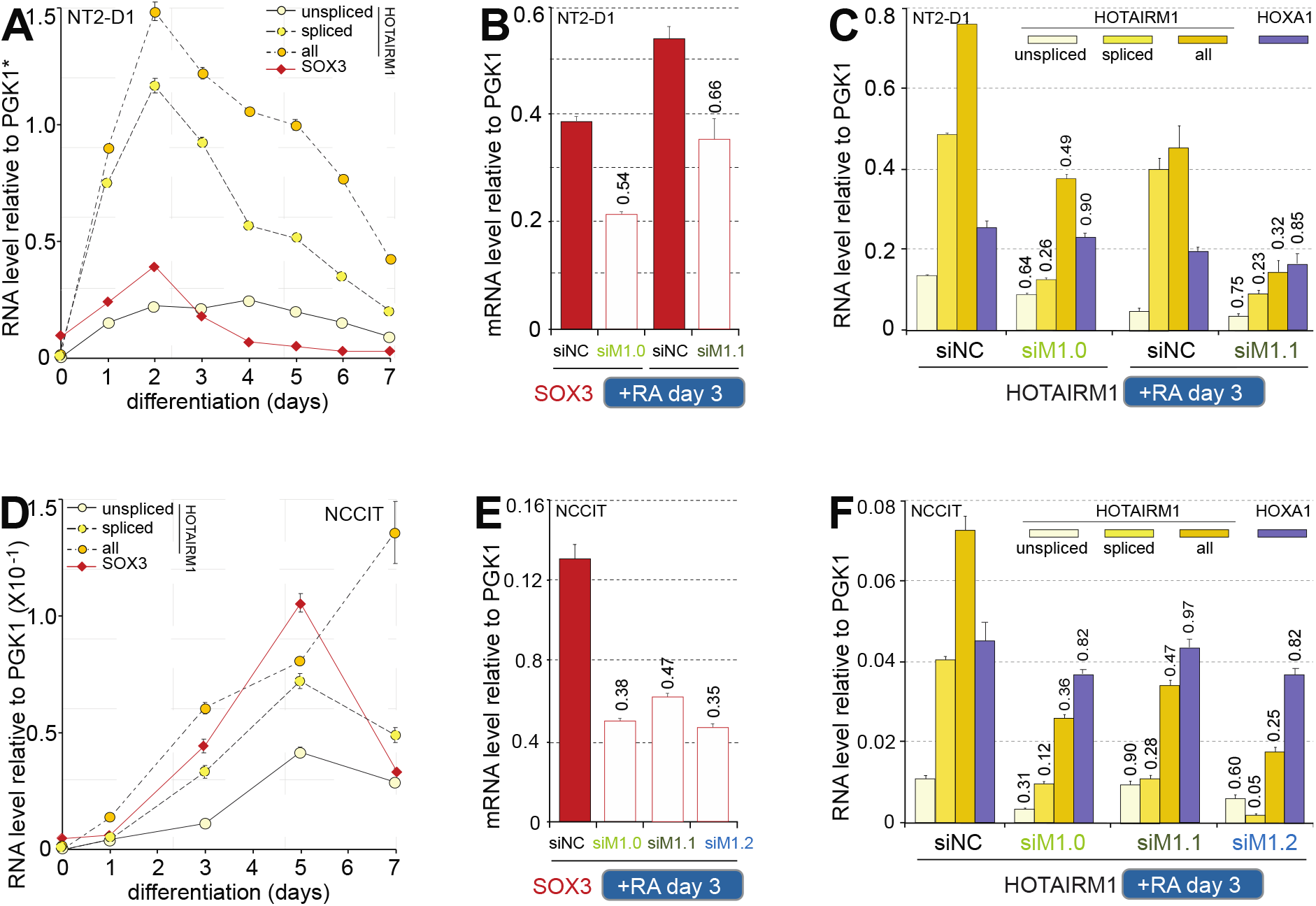
HOTAIRM1 is required for proper neuronal differentiation. (**A**) *SOX3* and *HOTAIRM1* expression during a differentiation time-course in NT2-D1. RNA levels were quantified by RT-qPCR using primers listed in Supplementary Table S1. Error bars are stdevs from 3 biological replicates, each with at least 3 measurements. \**SOX3* values are relative to PGK1 whereas those for HOTAIRM1 are relative to actin (10^−4^). (**B**,**C**) SOX3 (**B**) or HOTAIRM1 and HOXA1 (**C**) expression in RNAi knockdown samples of NT2-D1 treated 3 days with RA. Expression values are from at least 3 measurements each from 3 independent biological replicates. Error bars represent the (sem). Numbers above histogram bars indicate fold differences relative to corresponding siNC samples. (**D**) SOX3 and HOTAIRM1 expression during an NCCIT differentiation time-course. Expression levels were measured as in (**A**). A biological replicate extending to Day 14 post-RA is in Supplementary Figure S2A. (**E**,**F**) *SOX3* (**E**) or *HOTAIRM1* and *HOXA1* (**F**) expression in NCCIT RNAi knockdown samples treated 3 days with RA. Expression values are averages from at least 3 measurements and error bars are stdevs. Numbers above histogram bars are fold differences relative to corresponding siNC samples. A replicate of (**E**,**F**) collected 5 days post-RA is in Supplementary Figure S2B,C. A third biological replicate of Day 3 post-RA is given in Supplementary Figure S3E,F. Note that *HOXA1* values in (**C**,**F**) were divided by 10 to be on a scale comparable to HOTAIRM1.

We wondered if HOTAIRM1’s effect on *SOX3* was unique to the NT2-D1 cell line, and thus repeated the experiment in the human NCCIT differentiation model. NCCIT cells are developmentally pluripotent: they display EC and seminoma features and can be differentiated into derivatives of the three embryonic germ layers and extraembryonic lineages with RA [24]. In these cells, we also found SOX3 transiently and concurrently induced with HOTAIRM1, but with peaks delayed to 5 days post-RA treatment (Figure 3D). As in NT2-D1, NCCIT cells had was much lower *SOX3* expression when HOTAIRM1 was depleted by Day 3 (Figure 3E,F). This effect was recapitulated with 3 distinct siRNA targeting designs (siM1.0/1/2, Figure 2F), and in two replicates where cells were collected either at Day 3 during *SOX3* induction (Figure 3E,F) or at Day 5, when *SOX3* reaches peak expression (Supplementary Figure S2).

Given the transient nature of *SOX3* expression, and how rapidly *HOTAIRM1* is induced by RA to influence cell growth during differentiation, we asked whether depleting HOTAIRM1 curbs progress along the differentiation path or rather, accelerates it. We addressed this question by tracking *SOX3* expression levels throughout RA differentiation time-courses of control and HOTAIRM1-depleted cells (Supplementary Figure S3). In NT2-D1, *SOX3* induction appeared normal up to 48 hours post-RA treatment, but thereafter diverges from expected expression and fails to reach peak levels seen in control cells (Supplementary Figure S3A). These results were reproduced using two different RNAi targeting designs, and in separate biological replicates (Supplementary Figure S3; compare panel A to C). In NCCIT, *SOX3* induction deviated more rapidly upon RA induction, possibly because HOTAIRM1 was more potently knocked-down at these earlier time points compared to NT2-D1 (Supplementary Figure S3; compare panel F to B,D). Together, these results indicate that HOTAIRM1 presence is required for normal progression through the early stages of neuronal differentiation.

### HOTAIRM1 associates with the HOXA1 transcription factor

NT2-D1 cells have been extensively used to study *HOX* gene regulation as their response to RA recapitulates these genes’ collinear induction patterns in developing axial structures [25-28]. *HOX* genes are highly evolutionarily conserved, dating back to at least 550 Ma years ago [29]. They encode homeobox transcription factors that are master regulators of embryonic development with crucial roles in stem cell differentiation and cancers [30-32]. Several *HOX* genes have important functions during neurogenesis [33], including *HOXA1*, which is required for differentiation of stem cells into neurons in mouse [34,35]. Emerging information on the target genes and pathways controlled by HOXA1 reveals that it regulates very early steps of neuro-ectodermal differentiation in mouse ES cells [9,16,19,36].

We have shown that, like *HOTAIRM1, HOXA1* is highly and rapidly induced upon RA treatment of NT2-D1, whereas peak expression of the more distal *HOXA4/5/6* genes is reached later [7,12]. Though *HOXA1* and *HOTAIRM1* are transcribed from opposing DNA strands, they share a short promotor region that is highly conserved, especially between human and mouse (Figure 4A). *HOXA1*’s induction kinetics and expression levels are alike those of *HOTAIRM1* in the presence of RA (Figure 4B). Such analogous regulation suggests a functional link, and we postulated that HOTAIRM1 cooperates with the HOXA1 transcription factor as a ribonucleoprotein (RNP) complex.

**Figure 4.**
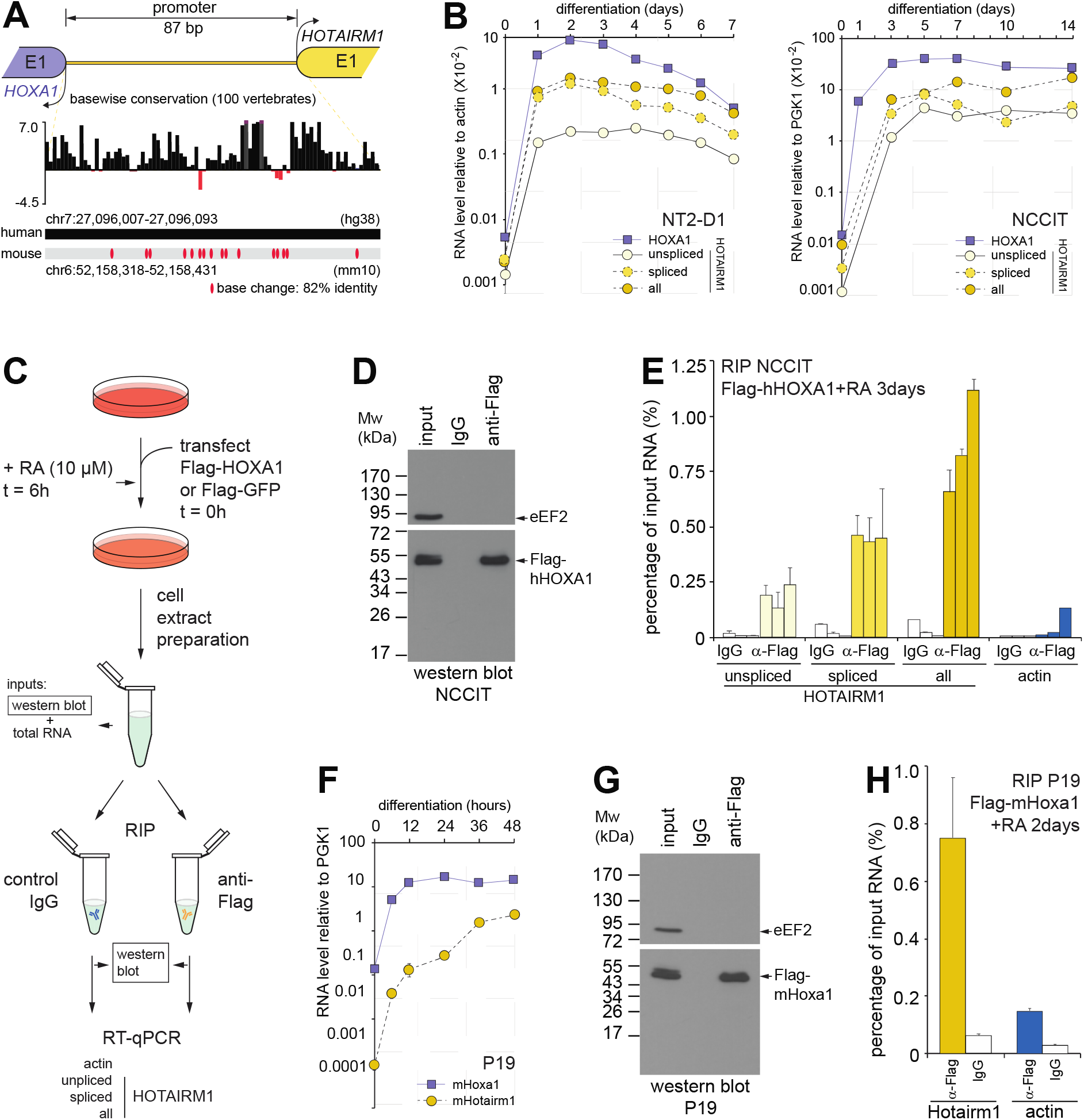
HOTAIRM1 associates with the HOXA1 transcription factor in different cell types and species. (**A**) Diagram of the highly conserved, shared promoter region from which *HOXA1* and *HOTAIRM1* are transcribed on opposite strands. Base-wise conservation across 100 vertebrates by phyloP and base change locations between human and mouse is shown below. (**B**) *HOXA1* and *HOTAIRM1* RA induction kinetics in two neuronal differentiation models. RNA levels measured during RA (10 μM) induction time-courses in NT2-D1 (*left*) or NCCIT (*right*) using RT-qPCR. (**C**) Outline of the RNA Immunoprecipitation (RIP) procedure. (**D**) Transfected Flag-hHOXA1 is specifically immunoprecipitated in RA-induced NCCIT cells. Western blot analysis of the input (1%) and RIP (5%) samples from one of the biological replicates analyzed in (**E**), with eukaryotic translation Elongation Factor 2 (eEF2) probed as a control for specificity. (**E**) HOXA1 preferentially binds spliced HOTAIRM1 in RA-induced NCCIT cells. RIPs are from 3 biological replicates quantified by RT-qPCR. (**F**) Induction of mouse *Hoxa1* and *Hotairm1* by RA (1 μM) in P19 cells. (**G**) Transfected Flag-mHoxA1 is specifically immunoprecipitated in RA-induced P19 cells. Western blot analysis of the input (1%) and RIP (5%) samples corresponding to the assay shown in (**H**). (**H**) Mouse HoxA1 preferentially binds mHotairm1 in RA-induced P19 cells. RT-qPCR measurements are from at least 3 PCRs with errors bars representing stdevs.

We tested this hypothesis by first performing RNA immunoprecipitations (RIPs) in RA-induced NCCIT cells transfected with a Flag-hHOXA1 construct (Figure 4C). We opted to transfect a tagged version of HOXA1 rather than probe the endogenous protein as, to our knowledge, no immunoprecipitation-grade anti-HOXA1 antibodies are available. Flag-hHOXA1 was specifically immunoprecipitated by the anti-Flag antibody as per the absence of endogenous eEF2 control protein and signal in control anti-IgG sample (Figure 4D). We found that Flag-hHOXA1 specifically associates with HOTAIRM1 in the corresponding sample and in two additional biological replicate RIPs (Figure 4E). HOTAIRM1 binding was specific to the HOXA1 protein as it did not bind to a Flag-GFP control (Supplementary Figure S4A,B). Notably, though HOXA1 bound both variants of HOTAIRM1, there was a marked preference for the spliced form.

We obtained similar results in RA-induced NT2-D1, despite higher cell death upon transfection in the cell model (Supplementary Figure S4C,D). Incidentally, we probed association of HOXA1 with HOTAIRM1 in HEK293T cells, where they are each constitutively expressed at high levels (Supplementary Figure S4E, *inset*). We found that HOXA1 associates with spliced HOTAIRM1 in these cells as well (Supplementary Figure S4E,F). This result was recapitulated in a separate biological replicate, and independent of UV cross-linking prior to the RIP (Supplementary Figure S4G,H). Binding of HOTAIRM1 to HOXA1 is therefore not unique to NCCIT cells or differentiating cell models. To determine if complex formation, as with these genes, is conserved across species, we used P19 cells to test if this interaction can be detected in the mouse (Figure 4F-H), as this EC line is also pluripotent and can be induced to differentiate into neuronal cells with RA [15]. We first verified induction of Hoxa1 and Hotairm1 by RA in a differentiation time-course (Figure 4F) before transfecting RA-induced P19 cells with Flag-mHoxa1 (Figure 4G). Again, we found that the tagged version of mouse Hoxa1 could be specifically co-immunoprecipitated with mouse Hotairm1 (Figure 4G,H). Together, our results show that HOXA1 inherently associates with HOTAIRM1 in vivo, regardless of cell type or species, and independently of the RA signaling pathway.

### HOXA1 and HOTAIRM1 modulate the state of the core pluripotency network

Given that HOXA1 and HOTAIRM1 can associate as an RNP, it stands to reason that they share some downstream functionality, and their gene targets must partly overlap. Amongst the known earliest targets of HOXA1 is the *NANOG* gene [16]. *NANOG* encodes a homeodomain-containing transcription factor essential for maintaining pluripotency along with OCT4 and SOX2, two other core pluripotency network components [37]. In the mouse, Nanog and Hoxa1 were shown to engage in a “double-negative gate” control mechanism [38]: while Nanog binds and represses the promoter and 3’ enhancer of Hoxa1 in undifferentiated cells, RA treatment prompts the expression and binding of Hoxa1 to *Nanog*’s promoter and autoregulatory enhancer (ARE), leading to down-regulation of the pluripotency gene (Figure 5A)[16,39]. At least in mESCs, Hoxa1 binds at the *Nanog* gene as early as 2 hours post-RA treatment, and coinciding with a decline in the gene’s transcriptional activity; this binding is dynamic – sometimes occurring very transiently in these early stages of neuro-ectoderm differentiation [16]. Using quantitative real-time PCR (RT-qPCR), we thus tracked the steady-state expression levels of *HOXA1* and *NANOG* throughout the early stages of neuronal differentiation and found the genes to be inversely correlated in both of our human cell models (Figure 5B). Also, in accordance with the shifted *HOTAIRM1* and *SOX3* expression kinetics true to our differentiation models (Figure 3A,D), we observed the rise of *HOXA1* and fall of *NANOG* to occur more rapidly in NT2-D1 as compared to NCCIT cells (Figure 5B, highlights).

**Figure 5.**
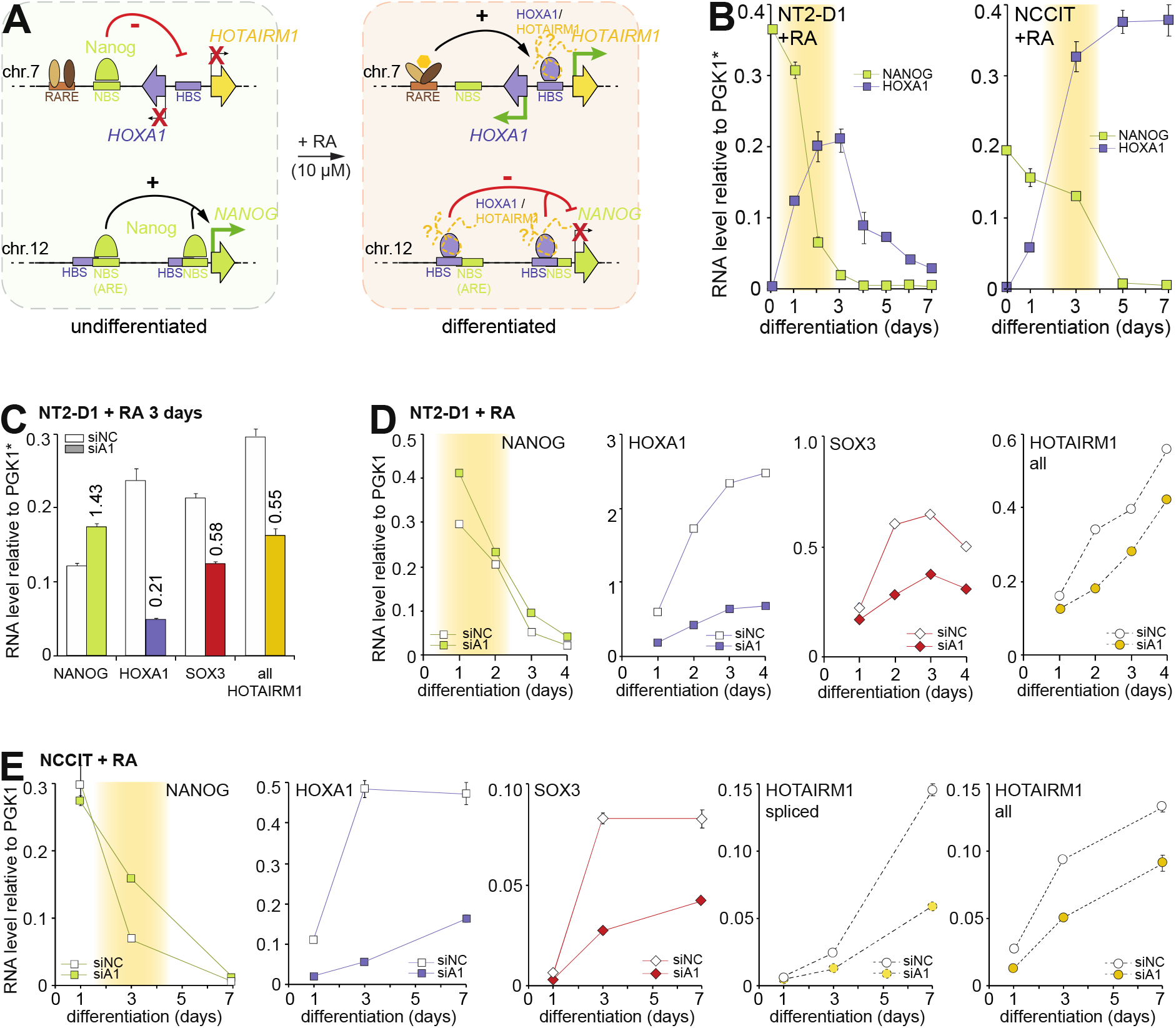
Depleting HOXA1 during differentiation delays NANOG downregulation. (**A**) NANOG and HOXA1 engage in positive auto- and negative cross-regulatory feedback mechanisms to control their gene expression. HOTAIRM1 (dashed) is proposed to work with HOXA1 as an RNP on the *NANOG* and *HOXA1/HOTAIRM1* loci in both undifferentiated and RA-induced cell states; RARE = Retinoic Acid Response Element, ARE = Autonomous Response Element, NBS = NANOG Binding Site, HBS = HOXA1 binding site. (**B**) Steady-state NANOG and HOXA1 mRNA levels throughout RA-induced differentiation of NT2-D1 (*left*) or NCCIT (*right*) cells. *Relative HOXA1 mRNA levels were divided by 10 to display on the same scale with NANOG. (**C**,**D**) HOXA1 knockdown delays *NANOG* downregulation, blunts *SOX3* upregulation, and impedes *HOTAIRM1* induction during NT2-D1 differentiation. HOXA1 was RNAi-depleted and cells were collected 3 days after RA treatment (**C**) or every 24 hours along a time-course (**D**). A similar differentiation time-course for control (siNC) and HOXA1 (siA1) knockdown cells in NCCIT is shown in (**E**). RNA levels were measured by RT-qPCR from at least 3 measurements, with error bars showing stdevs. Numbers above histogram bars are fold differences relative to levels in siNC samples.

We examined the impact of HOXA1 depletion on *NANOG* expression levels during NT2-D1 differentiation (Figure 5C,D). We detected approximately forty percent more NANOG transcripts 3 days post-RA treatment, at which time HOXA1 was considerably knocked down and *NANOG* was not yet wholly downregulated (Figure 5C). We then measured NANOG mRNA levels along a differentiation time-course of HOXA1-depeted NT2-D1 cells and found it consistently higher in HOXA1 knockdown samples, particularly at our earliest – 24h – timepoint (Figure 5D). Downregulation of *NANOG* was also delayed when HOXA1 was depleted in NCCIT cells (Figure 5E) at later timepoints, befitting this cell model’s temporal dynamics (Figure 5B, Figure 3).

Interestingly, *SOX3* expression was hampered by HOXA1 depletion in both NT2-D1 and NCCIT (Figure 5C-E), and to a similar extent as when HOTAIRM1 is knocked-down (Figure 3, Supplementary Figure S2B, Supplementary Figure S3A,C,E). Importantly, HOXA1 knockdown also reduced HOTAIRM1 levels in both cell differentiation models (Figure 5C-E). We repeated the experiment in NT2-D1 and verified that HOXA1 depletion indeed elicits lower HOTAIRM1 levels, since it does not significantly affect the expression of an unrelated housekeeping gene, *ACTB* (Supplementary Figure S5A). Furthermore, this effect was observed in our knockdowns using varying concentrations of siRNA, indicating that it is not an off-target effect of RNAi, but rather HOXA1 activating *HOTAIRM1* transcription (Supplementary Figure S5B).

Finding that HOTAIRM1 is reduced in HOXA1 knockdowns surmises that any gene expression changes cannot be attributed solely to the depleted transcription factor. We previously reported that HOTAIRM1 knockdown during NT2-D1 differentiation can – but does not always – lower *HOXA1* expression. RA induces approximately equal amounts of unspliced and spliced HOTAIRM1 transcripts in NT2-D1; whether or not HOXA1 levels are affected in HOTAIRM1 knockdown samples appears to require very efficient depletion of the unspliced form, which can only be achieved when using an shRNA against HOTAIRM1 [7]. In light of this, we verified that HOXA1 mRNA levels were not considerably affected in HOTAIRM1 siRNA knockdowns (Figure 2H, Figure 3C,F, Supplementary Figure S3B,D,F). We found that HOXA1 levels were only modestly and transiently affected when HOTAIRM1 was depleted at one timepoint, yet we observed higher NANOG levels along with lower SOX3 expression throughout NCCIT differentiation (Figure 6A,B). We obtained similar results when determining NANOG, HOXA1, and SOX3 levels early in differentiation of control and HOTAIRM1-depleted NT2-D1 cells (Figure 6C). These results together indicate that the impact of HOTAIRM1 depletion is not simply caused by sequent lowered HOXA1 levels, but rather indicates they share targets along a common regulatory axis.

**Figure 6.**
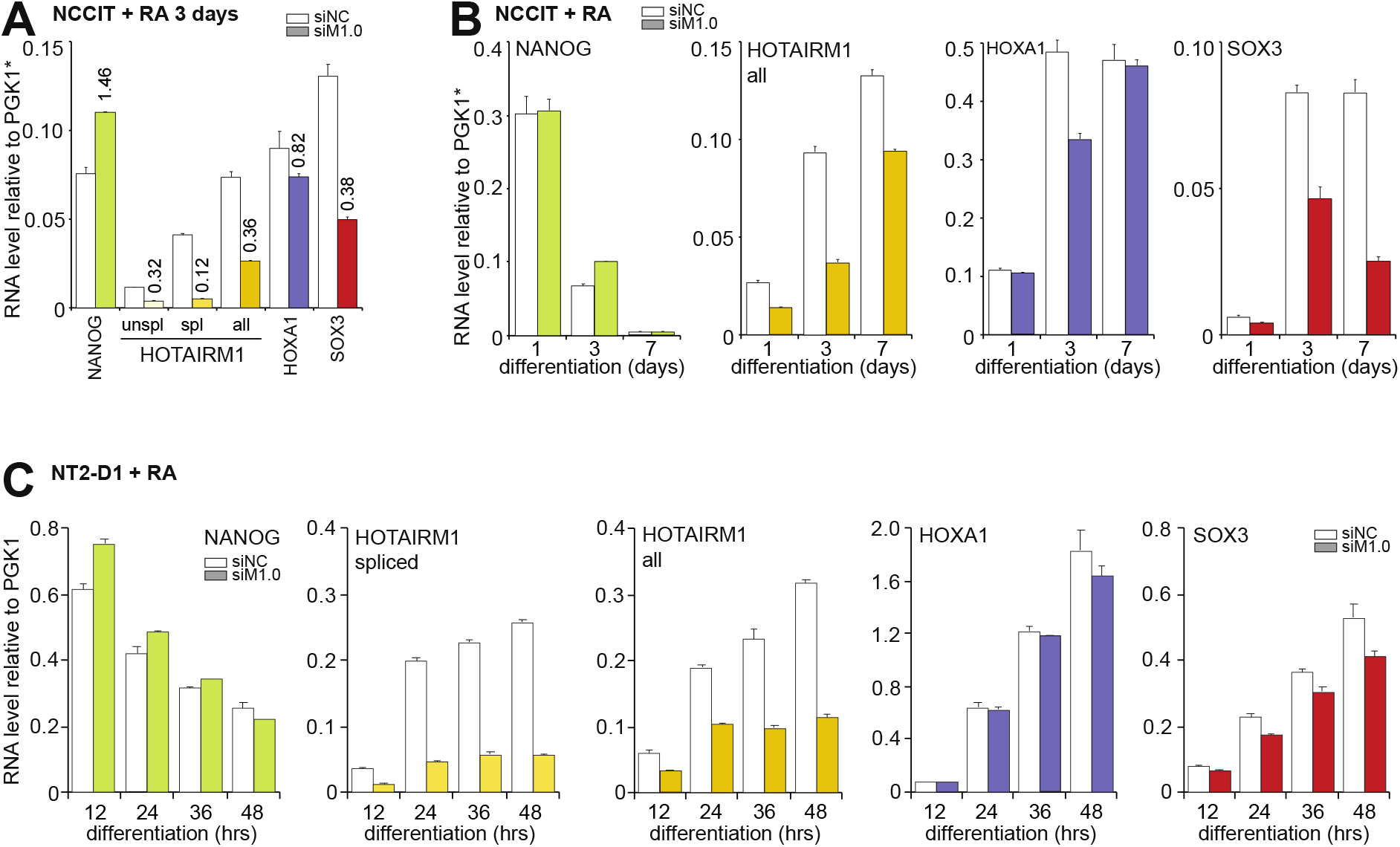
Depleting HOTAIRM1 during neuronal differentiation delays *NANOG* downregulation and *SOX3* upregulation. (**A**,**B**) RNAi depletion of HOTAIRM1 during RA-induced NCCIT differentiation. Control (siNC) or HOTAIRM1 (siM1.0)-depleted NCCIT cells were induced with RA and collected after 3 days (**A**), or at different times post RA-treatment (**B**). (**C**) Depleting HOTAIRM1 similarly affects *NANOG* and *SOX3* expression early in NT2-D1 differentiation. The experiment outlined in (**B**) was repeated in NT2-D1 cells, with samples collected at earlier timepoints. All RNA levels were measured by RT-qPCR from at least 3 measurements and error bars are stdevs. Numbers above histogram bars are fold differences relative to levels in siNC samples.

To further explore shared targets, we next examined whether the expression of other pluripotent factors is altered upon HOTAIRM1 depletion during differentiation. As mouse Hoxa1 has previously been reported to bind the *Sox2* locus rapidly upon RA stimulation in mESCs [16], we first profiled its expression in NT2-D1 and NCCIT during RA-induced differentiation and observed SOX2 levels progressively decrease upon RA treatment at a slightly faster rate in NT2-D1 compared to NCCIT, as expected (Figure 7A, compare panels *left* and *right*). In NT2-D1, we therefore tracked *SOX2* expression upon HOTAIRM1 depletion during early differentiation, when changes might be detected and under conditions where HOXA1 mRNA levels are not yet considerably affected. We found higher SOX2 levels when HOTAIRM1 is RNAi-depleted in both NT2-D1 and NCCIT (Figure 7B) and captured this effect in several biological replicates (Supplementary Figure S6). Interestingly, we also captured higher SOX2 levels when HOXA1 was knocked down in either of our differentiation models despite their inherent stochasticity (Figure 7C, Supplementary Figure S6C,D). However, levels of the third component in this core pluripotency network encoding OCT4, *POU5F1*, while modestly affected by HOXA1 depletion, remained unaffected when HOTAIRM1 was knocked-down during early differentiation (Figure 7D,E,F, Supplementary Figure S6).

**Figure 7.**
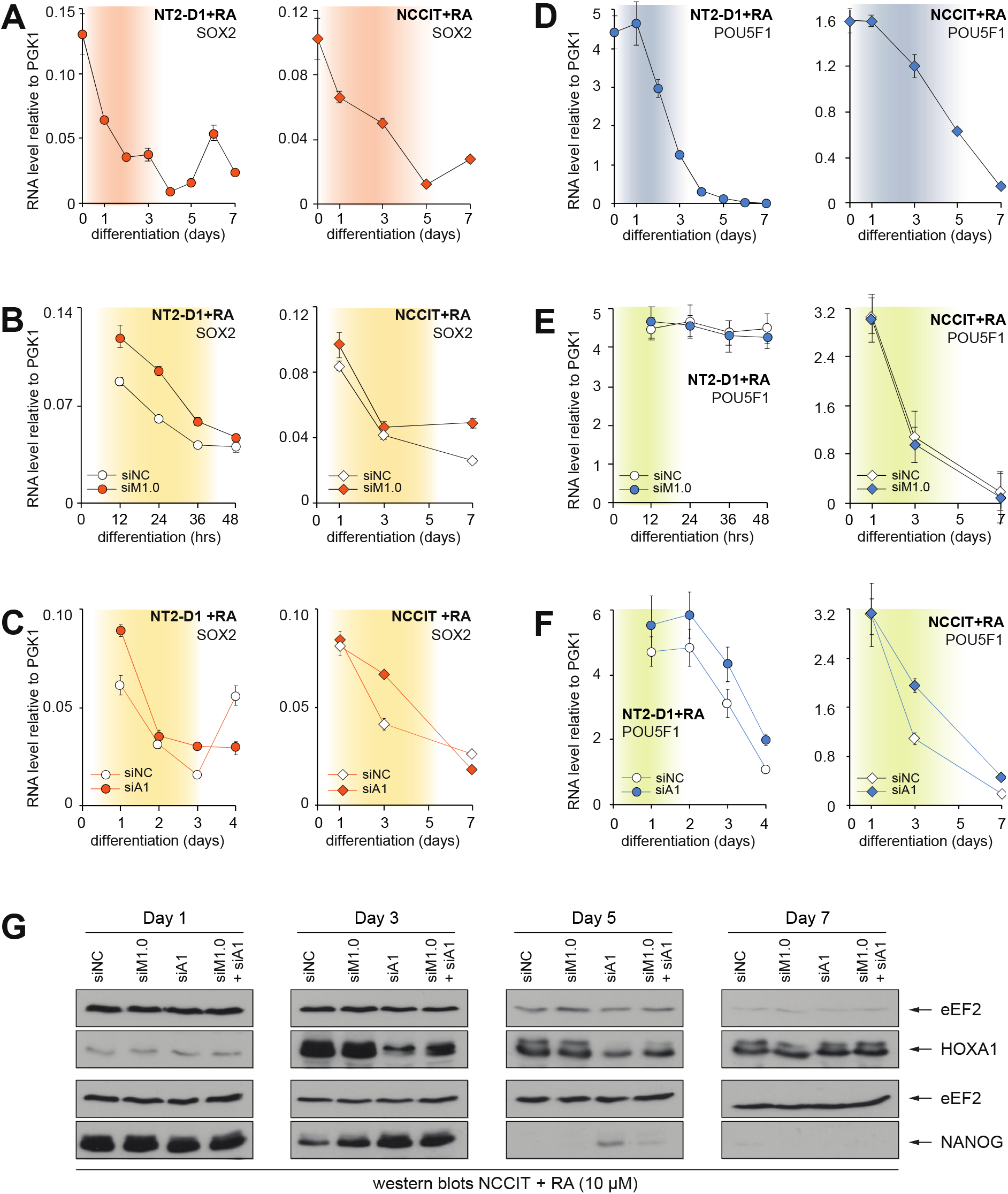
*SOX2* repression is more sensitive than *POU5F1* to HOTAIRM1 or HOXA1 depletion in early neuronal differentiation. (**A**) Steady-state SOX2 mRNA levels throughout RA-induced differentiation of NT2-D1 (*left*) or NCCIT (*right*) cells. (**B**) Depleting HOTAIRM1 during NT2-D1 (left) or NCCIT (right) differentiation curbs *SOX2* downregulation. (**C**) Same as in B, except upon RNAi-mediated *HOXA1* depletion. (**D**) Same as in (**A**), except that *POU5F1* expression was measured. (**E**) HOTAIRM1 depletion does not significantly affect *POU5F1* expression during either NT2-D1 (left) or NCCIT (right) differentiation. (**F**) *HOXA1* depletion modestly affects *POU5F1* expression during NT2-D1 (left) and NCCIT (right) differentiation. Knockdown levels of HOTAIRM1 and HOXA1 expression in (**B**) and (**E**) samples are shown in Figure 6B,C. HOXA1 knockdown levels and HOTAIRM1 expression in (**C**) and (**F**) samples are found in Figure 5D,E. All mRNA levels were measured by RT-qPCR and are from at least 3 measurements, with error bars representing stdevs. (**G**) Western blot analysis of HOXA1 and NANOG protein levels in either control (siNC), HOTAIRM1 (siM1) and/or HOXA1 (siA1) RNAi knockdown samples from a biological replicate RA differentiation time-course in NCCIT cells. The SOX2 and OCT4 protein levels and corresponding RNA levels measured by RT-qPCR and are presented in Supplementary Figure S6E,F.

### NANOG *and* SOX2 *are direct gene targets of the* HOTAIRM1-HOXA1 *regulatory axis*

The core pluripotent factor network’s highly interactive and self-modulated nature complicates the study of its regulation (Figure 8A, [16]). Indeed, the fact that components auto-regulate and influence each other, along with fluctuations inherent to biological replicates, is likely responsible for varying penetrance in our knockdowns. Regardless, we identified *NANOG* and *SOX2* as potential direct gene targets of HOXA1 and HOTAIRM1 in human cells, and thus sought to investigate if HOXA1 binds promoters and enhancers influencing these genes. In mouse, a Flag-tagged HoxA1 was reported to bind along the *HoxA* gene cluster as well as at enhancers neighboring *Nanog* and *Sox2* (Figure 8B)[16,19]. Though mechanistic details were not fully assessed, HoxA1 bound *Nanog*’s promoter and ARE, suggesting that it helps shut down *Nanog*’s transcription during differentiation.

**Figure 8.**
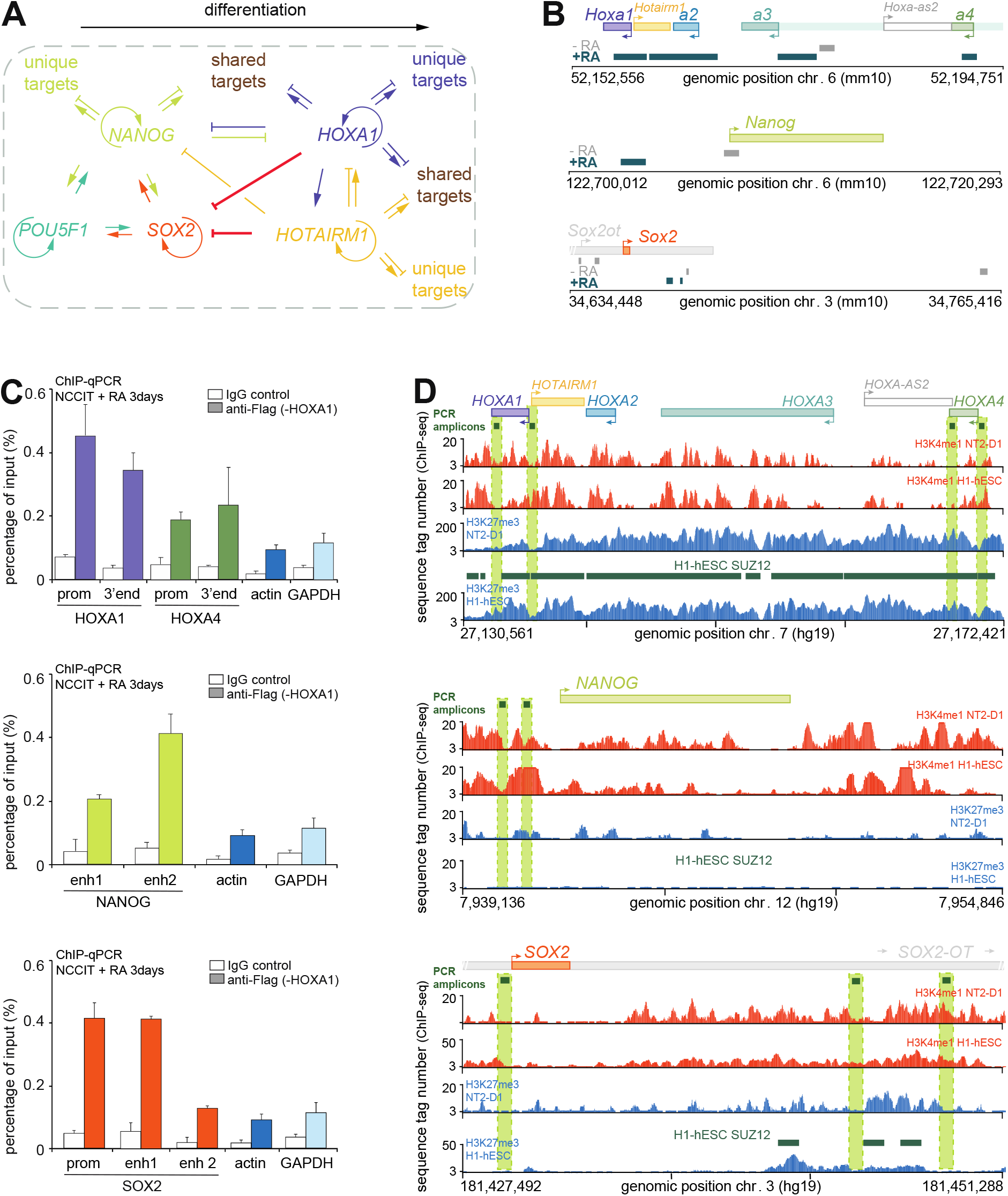
HOXA1 binding at *HOXA, NANOG*, and *SOX2* gene loci is conserved in human cells. (**A**) Summary schematic of the interaction network implicated with HOTAIRM1 and HOXA1. (**B**) Binding peaks of Flag-mHoxA1 in KH2 mES cells at 24h post-RA induction published in [19]. mHoxA1 binding is temporally dynamic, including at the mouse *Nanog* and *Sox2* promoters upon RA treatment in [16]. (**C**) ChIP-qPCR of Flag-hHOXA1 transfected in NCCIT treated 3 days with RA (10μM). The position of corresponding PCR amplicons is shown on the right in (**D**), dark green above ChIP-seq tracks, probing promoters (prom), 3’ ends, and enhancer (enh) regions. Negative control regions in the actin (*ACTB*) and *GAPDH* genes are repeated in each panel for comparison. Error bars are stdevs of at least 3 RT-qPCR measurements. A biological replicate is in Supplementary Figure S7. (**D**) Genomic position of PCR amplicons measured in (**C**). The H3K4me1 and H3K27me3 ChIP-seq tracks from undifferentiated NT2-D1 and H1-hESCs are presented below each locus to highlight proximity of probed regions to activating and repressive epigenetic chromatin marks.

We examined binding of human HOXA1 by transfecting a tagged version (Flag-hHOXA1) in RA-induced NCCIT cells followed by ChIP-qPCR, and found that, as in mice, HOXA1 binds the *HOXA1* and *HOXA4* promoters and gene bodies (Figure 8C, *top*). By BLAST sequence alignment, we could find no obvious homology to *Nanog*’s mouse ARE at the human *NANOG* locus, and thus probed two ENCODE candidate enhancers located upstream of the gene (E1590909/Enhancer1, E1590911/Enhancer2). While HOXA1 specifically bound the two regions, the association was stronger at Enhancer 2 (Figure 8C, *middle*), which incidentally corresponds to a region highly enriched for enhancer-associated mono-methylation at lysine 4 of histone H3 (H3K4me1) in both undifferentiated NT2-D1 and H1-hESCs (Figure 8D). Notably, Enhancer 2 is also marked by trimethylation at lysine 27 of histone H3 (H3K27me3) before NT2-D1 differentiation, a signature of bivalent chromatin, which could point to a potential silencing mechanism via Polycomb recruitment by the HOXA1/HOTAIRM1 RNP.

At the *Sox2* locus, tagged mouse HoxA1 was found to bind the *Sox2* promoter and downstream of the gene at two enhancer regions, but not at the distal control region (SRC) shown to be required for *Sox2* expression in mESCs [40]. While both remote binding sites correspond to enhancers as predicted by ENCODE [41], the site closest to *Sox2* displays the molecular signature of a proximal enhancer, is surrounded by ENCODE-predicted promoter signature sequences, and is immediately upstream of an annotated *Sox2ot* gene isoform. In contrast, the other Hoxa1 binding site, ENCODE-predicted enhancer beared a distal signature and was intergenic. Interestingly, this HoxA1 binding site overlapped with a previously validated enhancer shown to loop with the *Sox2* promoter in mESCs, but not in mouse embryonic fibroblasts (mEFs) [40].

We found that in human, transfected Flag-hHOXA1 also bound the human *SOX2* promoter in RA-induced NCCIT cells (Figure 8C, *bottom*). Importantly, we identified two regions homologous to the mouse enhancers and found HOXA1 binds at one of these sites (Figure 8C,D, *bottom panels*). Both human binding sites exhibited high sequence identity with their mouse counterparts, were intergenic, and overlapped with ENCODE-predicted distal enhancer elements. Moreover, these HOXA1-bound candidate distal enhancers either overlapped with or were adjacent to repressive H3K27me3 marks before NT2-D1 differentiation and in H1-hESC, suggesting that Polycomb recruitment to these regions may be involved also in *SOX2* repression at the onset of neurogenesis. Altogether, these results support the notion that HOXA1 and HOTAIRM1 share common regulatory targets along a control axis modulating the core pluripotency network early during neurogenesis.

## Discussion

In this study, we show that the HOTAIRM1 lncRNA is required for proper progression through the early stages of neuronal differentiation. We discovered that it associates with the HOXA1 transcription factor, likewise necessary for neuronal differentiation. Binding of HOTAIRM1 and HOXA1 in an RNP complex was observed regardless of cell type, RA signaling, experimental parameters, as well as in mouse (Figure 4, Supplementary Figure S4). *HOTAIRM1* and *HOXA1* are rapidly induced by RA from a shared promoter region (Figure 4A); this genomic configuration is likely meant to coincide their expression to coordinate their roles. Interestingly, *HOXA13* and *HOTTIP* share a similar antisense arrangement at the opposite end of the *HOXA* cluster, with both eliciting transcriptional control at the *BMP7* gene promoter in gastric cancer cells [42]. To our knowledge, we are first to report of a physical coupling between a *HOX* transcription factor and neighboring lncRNA that function in the context of cell differentiation. These findings signify that functional coupling might exist for other such RNP complexes along the *HOXA* gene cluster, and possibly throughout the genome.

Further supporting the premise that linear organization underlies function at the *HOXA* cluster is the high evolutionary conservation observed in terms of synteny and sequence [6,43]. Beyond the notion that evolutionary conservation usually reflects important functional properties, the conserved *HOTAIRM1* and *HOXA* expression domains, their co-activation, and co-localization during development in amphioxus and xenopus [6] indicates conservation of their functional co-operation as well. HOTAIRM1 likely plays a role in neuronal differentiation across species, as interfering with *HOTAIRM1* expression during X. tropicalis development results in embryonic posteriorization with a severe headless phenotype [6].

Because *HOTAIRM1* and *HOXA1* are such evolutionarily conserved genes, we postulated that they together regulate control networks fundamental to development. Since we observed that HOTAIRM1 depletion prevents proliferation (Figure 2, Supplementary Figure S1) earlier than prevent neuronal marker *SOX3* induction (Figure 3, Supplementary Figure S2), we viewed the *SOX3* defect as to be a consequence of differentiation malfunction and considered possible deregulation of transcription factors central to maintaining pluripotency. Indeed, *NANOG* and *SOX2* failed to properly downregulate when either HOTAIRM1 or HOXA1 are depleted (Figs. 5-7, Supplementary Figure S6). Although we have not yet defined the mechanism by which HOTAIRM1 regulates these genes, it is within reason to speculate it is partly through co-recruitment of HOXA1 at promoters and neighboring enhancers (Figure 8C,D). We show that the HOXA1 transcription factor is recruited along the human *NANOG* and *SOX2* loci (Figure 8B), corresponding to similar sites in mouse [19]. Considering our previous finding that HOTAIRM1 binds components of the polycomb repressive complex 2 (PRC2) [7], we propose that the lncRNA might facilitate polycomb recruitment to these sites via HOXA1 binding to silence them during differentiation.

In addition, we show human HOXA1 binding at its own promoter region shared with HOTAIRM1, and to HOXA4 as reported in mESCs (Figure 8B,C,D, *top panels*; [16,19]). This result, along with finding that HOXA1 depletion consistently leads to lower HOTAIRM1 levels (Figure 5C-E, Supplementary Figure S6C,D) indicates that it likely positively regulates HOTAIRM1 expression during differentiation, in addition to auto-upregulating its own transcription. Together with our previous report that unspliced HOTAIRM1 can activate *HOXA1* expression by binding and recruiting MLL/SET complexes at the gene [7], these results point to HOXA1 and HOTAIRM1 engaging into a positive feedback mechanism where their expression enhances each other’s in early neuronal differentiation. This reciprocal feedback loop might be a conserved feature of the gene pair, given that in myeloid cells, either genes’ ectopic overexpression or depletion enhances or downregulates the other, respectively [44].

Since knockdown of HOXA1 consistently leads to lower HOTAIRM1 levels, we cannot precisely assign HOXA1’s contribution to the gene expression defects observed in our experiments. Nonetheless HOXA1 does appear to play a role given that gene expression defects are more pronounced in HOXA1 knockdowns while HOTAIRM1 transcript considerably remains, relative to siRNA-mediated HOTAIRM1 depletion. In contrast, as HOXA1 levels are not substantially affected under the HOTAIRM1 knockdown conditions used in this study (Figs. 2H, 3C, 3F, 6), we can reasonably implicate HOTAIRM1 in *NANOG* and *SOX2* silencing early in differentiation. While we have explored the conservation of Hoxa1 and Hotairm1’s physical association in mouse, we have yet to evaluate if their gene expression effects are conserved across species.

Though molecular details on HOTAIRM1’s regulation remain to be elucidated, at the *SOX2* locus, HOTAIRM1 might recruit polycomb complexes via HOXA1 to dampen transcriptional activity at promoters and enhancers. We identified two regions at the human *SOX2* locus that are homologous to distal mouse enhancers bound by HoxA1 in RA-treated mESCs [40]. Both of these candidate human enhancer regions – Enhancer 1 and Enhancer 2 – exhibited high sequence identity to their mouse counterparts, were intergenic, and overlapped with ENCODE-predicted distal enhancer sequences. Human Enhancer 2 corresponds to mouse SRR18, shown to loop with the *Sox2* promoter specifically in mESCs [40]. Though we only detected binding to Enhancer 1 at 3 days post-RA treatment (Figure 8C, Supplementary Figure S7A), we suspect that transfected Flag-HOXA1 could bind Enhancer 2 at an earlier timepoint, envisioning conservation of Hoxa1’s dynamic chromatin binding behavior seen in mouse [16]. In fact, mouse SRR18 is not required for *Sox2* transcription in mESCs, raising the possibility that this enhancer-promoter loop contributes more to *Sox2* silencing via an HOXA1/HOTAIRM1 RNP; it will be interesting to determine whether this represents the underlying mechanism switching off *SOX2* during neuronal differentiation.

Finally, one might ponder why we were able to capture gene expression effects on the core pluripotent factor network for *NANOG* and *SOX2*, but not as consistently for *POU5F1* (Figure 8A). *POU5F1* likewise upregulates itself and the other two components of the network; however, RT-qPCR analyses show that *POU5F1* transcript levels are much higher – at least ten times greater – than those of *NANOG* or *SOX2* in our cell models (Figure 7D,E,F), potentially dampening sensitivity to HOTAIRM1 or HOXA1 depletion. The continuous autoactivation of *POU5F1* and its impact on *NANOG* and *SOX2* transcription might also contribute to the apparent stochasticity of our knockdown systems. Deciphering the role of lncRNAs in general is a challenging task, not only because they tend to modulate pathways rather than act as molecular switches, but also as they impinge on complex pathways. Nevertheless, our study points to the existence of a complex, yet conserved, HOTAIRM1-HOXA1 regulatory axis essential for proper neuronal differentiation. It will be compelling to explore which other genes are directly targeted by HOTAIRM1 and/or HOXA1, and molecularly dissect the various mechanisms by which they exert influence, both during differentiation and in cancers where they are aberrantly expressed.

## Data Availability

The ChIP-seq peak tracks for Flag-mHoxa1-expressing KH2 mESCs were published in [19] and can be found at https://genome.cshlp.org/content/27/9/1501/suppl/DC1. The NT2-D1 and H1-hESC ChIP-seq datasets for H3K4me1 (wgEncodeEH000917, wgEncodeEH000106) and H3K27me3 (wgEncodeEH000908, wgEncodeEH000074) can be found on UCSC genome browser (http://genome.ucsc.edu).

## Accession Numbers

The accession numbers for NT2-D1 and H1-hESC ChIP-seq datasets, respectively, for H3K4me1 are wgEncodeEH000917 and wgEncodeEH000106, and for H3K27me3 are wgEncodeEH000908 and wgEncodeEH000074.

## Supplementary Data

Supplementary Data are available at NAR online.

## Acknowledgements

We thank members of our laboratory for insightful discussions. We specifically thank D. Paquette, M. Bellefeuille, C.Chin Sang, and A. Groff for their technical assistance.

## Author Contributions

J.D. conceptualized the study. J.D. and D.S. designed the experiments. D.S. conducted most experiments: J.S. assisted with the live cell imaging and S.C. handled bioinformatics data. J.D. and D.S. wrote the manuscript.

