## Supplemental Material for "A conserved *HOTAIRM1-HOXA1* regulatory axis coordinates early neuronal differentiation"

#### List of supplemental material relating to the main manuscript:

Supplemental Materials and methods: This section completes the "Materials and methods" section of the main text by including important technical details.

Supplemental Figures and Tables:

Supplemental Table 1: A list of primer sequences used for real-time quantitative PCR analysis in this study.

Supplemental Figure 1: This figure presents the results of a biological replicate of an experiment included in Figure 2.

Supplemental Figure 2: This figure presents the results of biological replicates of experiments included in Figure 3D,E,F.

Supplemental Figure 3: This figure relates to Figure 3. It features RA induction time-courses of HOTAIRM1 knockdown cells (NT2-D1 and NCCIT) to track SOX3 reduction through time.

Supplemental Figure 4: This figure relates to Figure 4. It features RIPs (RNA immunoprecipitations) in NCCIT, NT2-D1, and HEK293T cells that complement RIPs in Figure 4.

Supplemental Figure 5: This figure relates to Figure 5. It features biological replicates of HOXA1 (siA1) and control (siNC) RNA knockdown samples confirming that HOXA1 depletion specifically impedes HOTAIRM1 induction.

Supplemental Figure 6: This figure relates to Figure 7 and features biological replicates of control (siNC) and HOTAIRM1 (siM1.0) knockdowns over an extended time-course in NT2-D1 (*A*,*C*) or after 3 days of RA treatment in NCCIT (*B*,*D*) to confirm knockdown effects on *SOX2* and *POU5F1* expression. It also features a separate biological replicate time-course of control (siNC), HOTAIRM1 (siM1.0), HOXA1 (siA1), HOTAIRM1+HOXA1 (siM1.0+siA1) knockdowns in NCCIT cells (*E*,*F*).

Supplemental Figure 7: This figure relates to Figure 8 and features a biological replicate for the ChIP-qPCR analysis of transfected Flag-hHOXA1 in NCCIT cells treated with RA (10 µM) for 3 days.

#### Supplemental Materials and Methods

##### *Cell culture*

HEK293T cells were kindly provided by Dr. J. Teodoro (McGill, ATCC; CRL-3216). They are human embryonic kidney cells and were cultured in DMEM supplemented with 10% FBS (Gibco™). The cells were grown at 37°C in 5% $CO_2$ atmosphere.

##### *RNA immunoprecipitation (RIP) with UV cross-linked cell samples (CLIP)*

Cells were grown to 70% confluency prior to UV-crosslinking and washed once with 1X PBS. Cells were kept on ice while subjected to 100 mJ/cm$^2$ UV irradiation at 254 nm using a Hoefer™ UVC 500 Ultraviolet Crosslinker. After replacing with fresh, cold 1X PBS, the cells were scraped into solution and pelleted by centrifugation. Following a wash with cold 1X PBS, cells were lysed in equal volume lysis buffer for CLIP.

### Supplemental Figures and Tables

Segal_Supplemental_Table1

**Supplemental Table 1.** DNA sequence of primers used for quantitative real-time reverse transcription polymerase reaction (R T-qPCR).

| Specie | Gene name or region | Primer name | Primer sequence (5'-3') |
|---|---|---|---|
| human | *HOTAIRM1* | Exon3 Fwd1 | TAGTTATTGACCTGGAGACTGGTAGC |
| human | *HOTAIRM1* | Exon3 Rev1 | TCAGTGCACAGGTTCAAGCC |
| human | *HOTAIRM1* | Exon2 Fwd | AGGGAAGGTAGGGAGCAAACCTATG |
| human | *HOTAIRM1* | Exon3 Rev3 | GTTGATGGGTTCAGGCAAAACAGAC |
| human | *HOTAIRM1* | Intron1 Fwd | CTGGAGCTGGTCTCTTTCAACG |
| human | *HOTAIRM1* | Intron1 Rev | CCTTTCTAGCATAAGAGCCC |
| human | *GAPDH* | GAPDH Fwd | CCCAGCAAGAGCACAAGAGG |
| human | *GAPDH* | GAPDH Rev | TGGTACATGACAAGGTGCGG |
| human | *PGK1* | PGK1 Fwd | CTGTGCCAAATGGAACACGG |
| human | *PGK1* | PGK1 Rev | ATTGCTGAGAGCATCCACCC |
| human | *ACTIN* | ACTIN Fwd | CCTGGCACCCAGCACAATGAAG |
| human | *ACTIN* | ACTIN Rev | AAGTCATAGTCCGCCTAGAAGC |
| human | *SOX3* | SOX3 Fwd | TTGTAGGCTGGGAATCGCTG |
| human | *SOX3* | SOX3 Rev | TAGGCGTTGCAGTTCTCCAG |
| human | *HOXA1 (3'end)* | HOXA1 Fwd | TCCTAAGACCCGTAAACTCTGC |
| human | *HOXA1 (3'end)* | HOXA1 Rev | GCATGTCGCACAATGTTTGATG |
| human | *HOXA4 (3'end)* | HOXA4 Fwd | CAGAAGGGGACAACAGTATCTC |
| human | *HOXA4 (3'end)* | HOXA4 Rev | CATTAAGGCAGCTCATCCAAGC |
| human | *NANOG* | NANOG Fwd | GTCTCGTATTTGCTGCATCGT |
| human | *NANOG* | NANOG Rev | AACACTCGGTGAAATCAGGGT |
| human | *SOX2* | SOX2 Fwd | GTGAGCGCCCTGCAGTACAA |
| human | *SOX2* | SOX2 Rev | GCGAGTAGGACATGCTGTAGGTG |
| human | *POU5F1* | POU5F1 Fwd | AAATTCTCCAGGTTGCCTCT |
| human | *POU5F1* | POU5F1 Rev | GAAGGTATTCAGCCAAACGA |
| human | *HOXA1prom* | HOXA1 prom Fwd | CTGCTAAGTATGGGGTATTCC |
| human | *HOXA1prom* | HOXA1 prom Rev | TGGAGGAAGTGAGAAAGTTGG |
| human | *HOXA4prom* | HOXA4 prom Fwd | TTCAGGGGTTCTAGGCTAAC |
| human | *HOXA4prom* | HOXA4 prom Rev | GGTTGAGAAAATGTGTCTGG |
| human | *NANOGenh1* | NANOG enh1 Fwd | GGTTAAACAGAGCTTTCCCCCA |
| human | *NANOGenh1* | NANOG enh1 Rev | GAAATGAAGGCAAACGGCAGG |
| human | *NANOGenh2* | NANOG enh2 Fwd | GTTAGTGCTGGAACCCCACTCT |
| human | *NANOGenh2* | NANOG enh2 Rev | AAGACTACTCCGTGCCCATCT |
| human | *SOX2prom* | SOX2 prom Fwd | GCGCTGATTGGTCGCTAGAA |
| human | *SOX2prom* | SOX2 prom Rev | TCTCTGCCTTGACAACTCCTGA |
| human | *SOX2enh1* | SOX2 enh1 Fwd | TTCCATATTTGCGCCTCCCG |
| human | *SOX2enh1* | SOX2 enh1 Rev | ACCGCGAGCTTTTTCGTTTC |
| human | *SOX2enh2* | SOX2 enh2 Fwd | CTTTCGGATTCCATAGACAATCTCC |
| human | *SOX2enh2* | SOX2 enh2 Rev | CAAGTAGTCTGAAAGTTATGGGAAC |
| mouse | *Hoxa1* | Hoxa1 Fwd | CAGCGCAGACCTTTGACTGG |
| mouse | *Hoxa1* | Hoxa1 Rev | GCGCTCGTGTAAGGTACTTGT |
| mouse | *Hotairm1* | Hotairm1 Fwd | AGCTGGGAGATTAATCAACC |
| mouse | *Hotairm1* | Hotairm1 Rev | GAGTTCCCTTACCAAGCTGC |
| mouse | *Actin* | actin Fwd | TACTCTGTGTGGATCGGTGG |
| mouse | *Actin* | actin Rev | ACGCAGCTCAGTAACAGTCC |

* HOXA1 (3'end), HOXA4 (3'end), actin, and GAPDH primers were used for both gene expression and ChIP measurements.

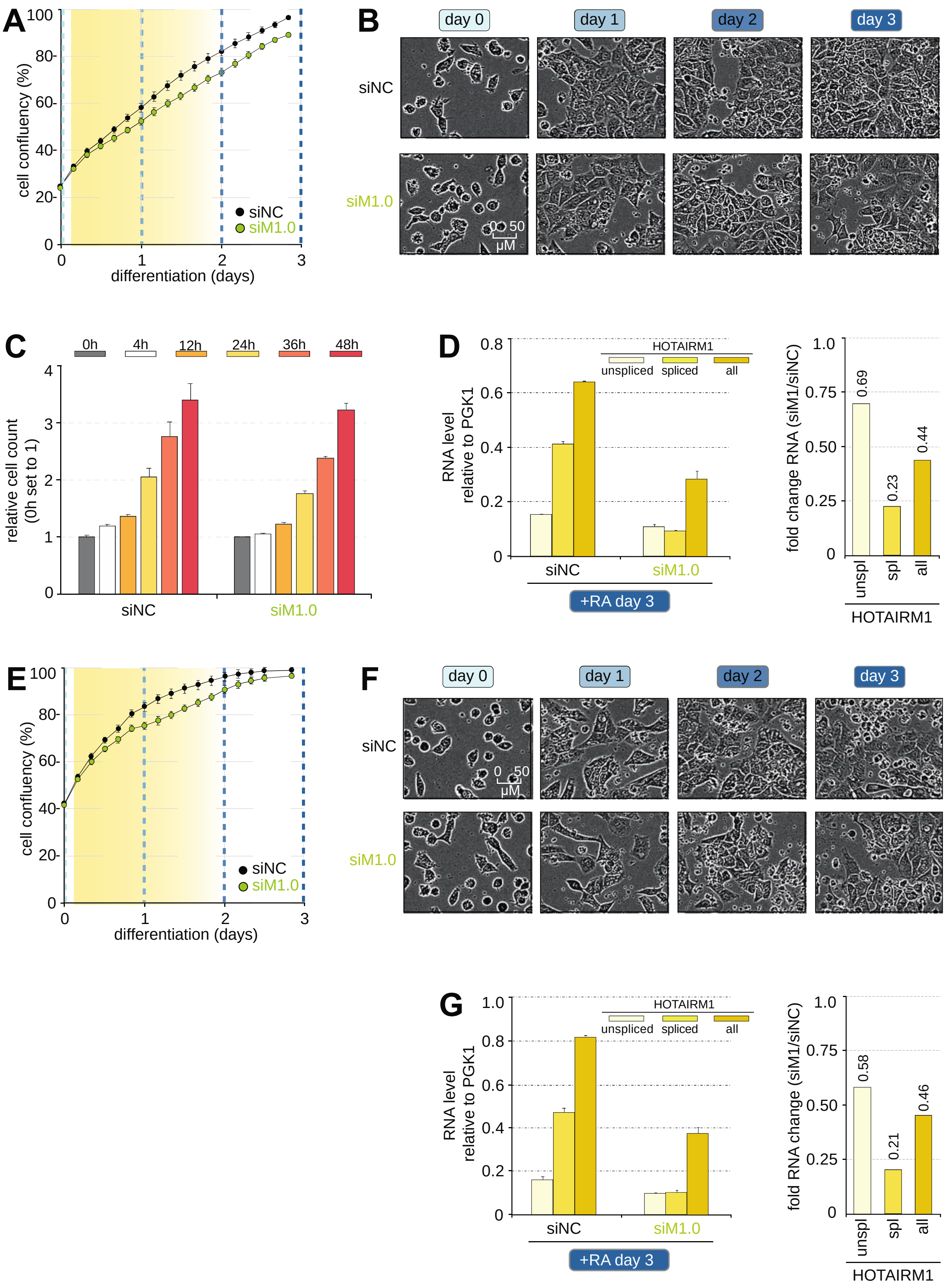
A
cell confluency (%)
differentiation (days)
siNC
siM1.0
B
day 0
day 1
day 2
day 3
siNC
siM1.0
0 50 μM
C
0h
4h
12h
24h
36h
48h
relative cell count (0h set to 1)
siNC
siM1.0
D
HOTAIRM1
unspliced
spliced
all
RNA level relative to PGK1
siNC
siM1.0
+RA day 3
fold change RNA (siM1/siNC)
0.69
0.23
0.44
unspl
spl
all
HOTAIRM1
E
cell confluency (%)
differentiation (days)
siNC
siM1.0
F
day 0
day 1
day 2
day 3
siNC
siM1.0
0 50 μM
G
HOTAIRM1
unspliced
spliced
all
RNA level relative to PGK1
siNC
siM1.0
+RA day 3
fold RNA change (siM1/siNC)
0.58
0.21
0.46
unspl
spl
all
HOTAIRM1

**Supplemental Figure 1.** Cell proliferation is hindered in two additional biological replicates when HOTAIRM1 is RNAi-depleted during RA-induced neuronal differentiation. (*A*) Confluency changes during RA differentiation of a second biological replicate of HOTAIRM1-depleted NT2-D1 cells as captured by live cell imaging with an Incucyte® Zoom analysis system. Data was collected as described in Fig. 2B and C. (*B*) Representative phase contrast microscopy images of NT2-D1 cells throughout the differentiation time-course monitored in (*A*). (*C*) Average cell number manually curated from Incucyte® images of 12 culture dish regions. Values are relative to the counted 0h seeding number, set to 1. Error bars show the standard error of the mean across the 12 culture dish regions. (*D*) Steady-state expression levels of HOTAIRM1 variants in RNAi knockdown samples treated with RA for 3 days and monitored in (*A*). Expression levels are shown relative to the *PGK1* housekeeping gene. Error bars are the standard deviations between at least 3 RT-qPCR measurements. Numbers above histogram bars on the left indicate fold differences. (*E*) Confluency changes during RA differentiation of a third biological replicate of HOTAIRM1-depleted NT2-D1 cells as described above in *A*. (*F*) Representative phase contrast microscopy images of NT2-D1 cells throughout the differentiation time-course monitored in (*E*). (*G*) Steady-state expression levels of HOTAIRM1 variants in RNAi knockdown samples treated with RA for 3 days and monitored in (*E*). Expression levels are shown relative to the *PGK1* housekeeping gene. Error bars are the standard deviations between at least 3 RT-qPCR measurements. Numbers above histogram bars on the left indicate fold differences. Yellow shading in (*A*,*E*) highlights growth delay when HOTAIRM1 is depleted.

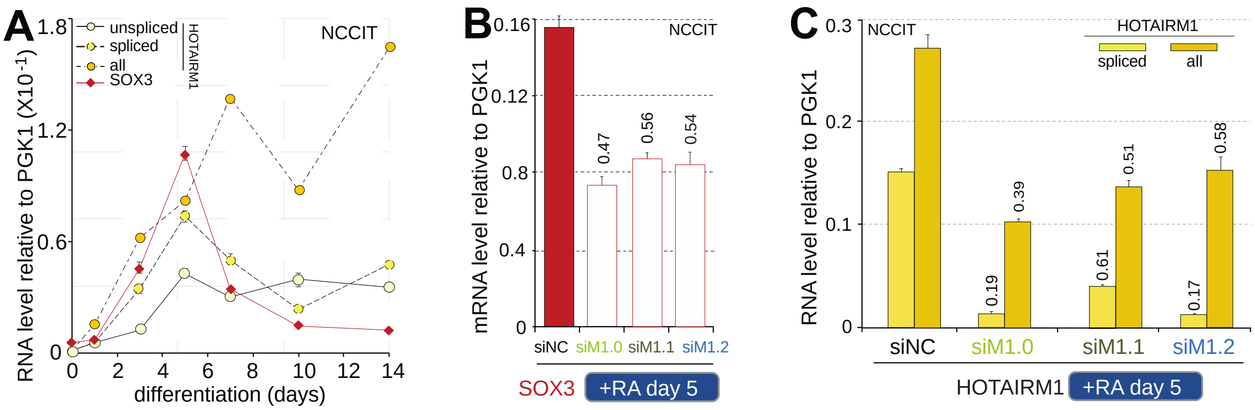


**Supplemental Figure 2.** HOTAIRM1 is required for proper RA-induced neuronal differentiation. (*A*) Biological replicate of the *SOX3* and HOTAIRM1 induction time-course in NCCIT as shown in Fig. 3D. Expression levels were measured as described in Fig. 3A. Error bars are standard deviations from at least 3 RT-qPCR measurements.

(*B*,*C*) Replicate of Fig. 3E,F at 5 days post RA-induction showing SOX3 (*B*) or HOTAIRM1 (*C*) expression in NCCIT RNAi knockdown samples. A biological replicate of this experiment is a component of the knockdown time-course presented in Supplemental Fig. 3E,F. Expression values are averages from at least 3 RT-qPCR measurements and error bars denote standard deviations.

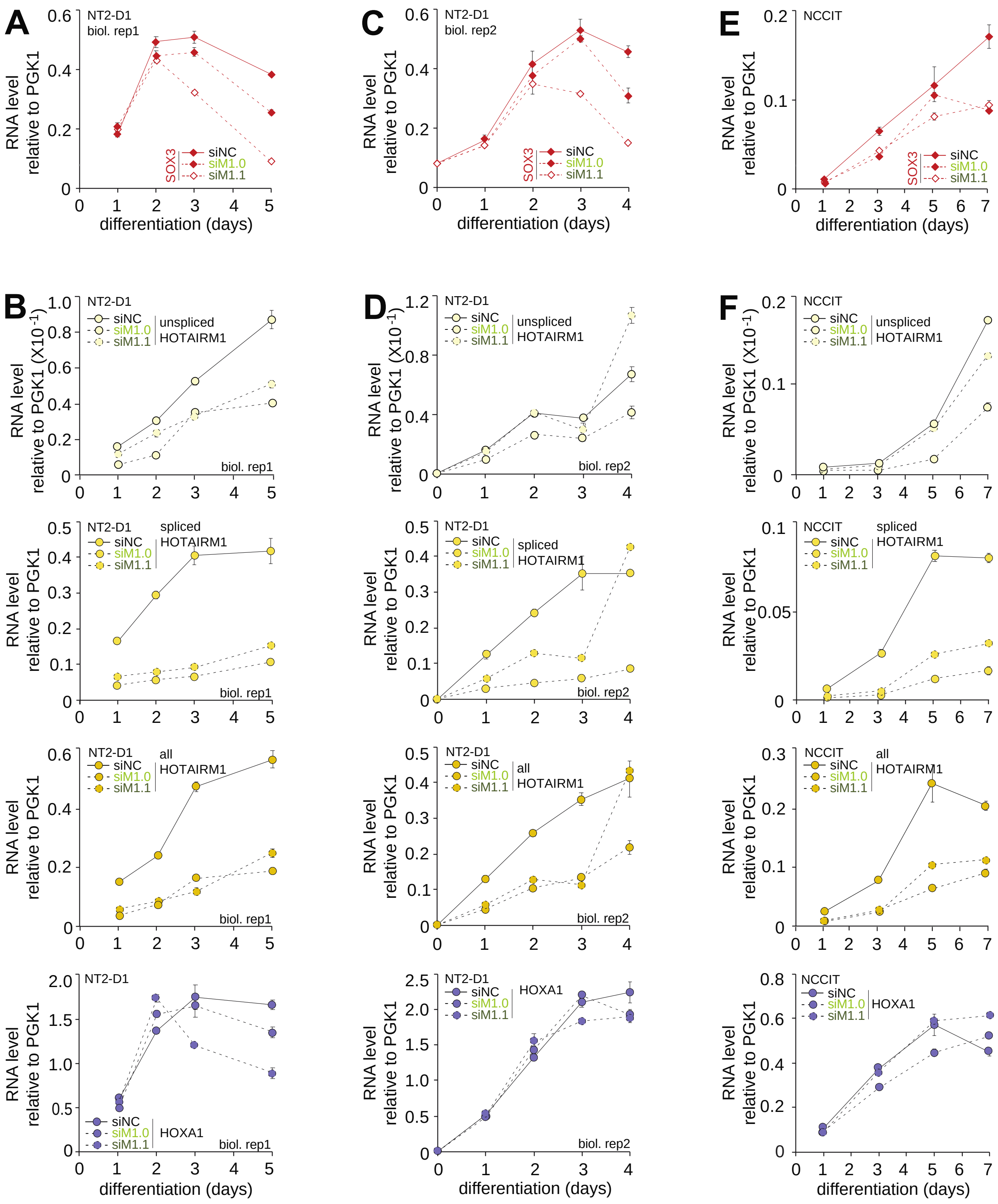
A
NT2-D1
biol. rep1
RNA level relative to PGK1
SOX3
siNC
siM1.0
siM1.1
differentiation (days)
C
NT2-D1
biol. rep2
E
NCCIT
B
unspliced HOTAIRM1
RNA level relative to PGK1 (X10 -1)
spliced HOTAIRM1
all HOTAIRM1
HOXA1
D
F

**Supplemental Figure 3.** Time-course of *SOX3* expression in RA-induced HOTAIRM1 knockdown cells. (*A*,*B*) Steady-state SOX3 (*A*) or HOTAIRM1 and HOXA1 (*B*) RNA levels in control (siNC) or HOTAIRM1 (siM1.0, siM1.1) NT2-D1 knockdown cells induced with RA (10 µM) were quantified by RT-qPCR as outlined in Materials and methods. Error bars are standard deviations from at least 3 measurements. (*C*,*D*) Biological replicate of panels *A*,*B* except that the time-course extends to only Day 4 post RA-induction. (*E*,*F*) Time-course of SOX3 (*E*) or HOTAIRM1 and HOXA1 (*F*) expression in RA-induced, NCCIT HOTAIRM1 knockdown NCCIT cells. Steady-state RNA levels are as described in *A*,*B*.

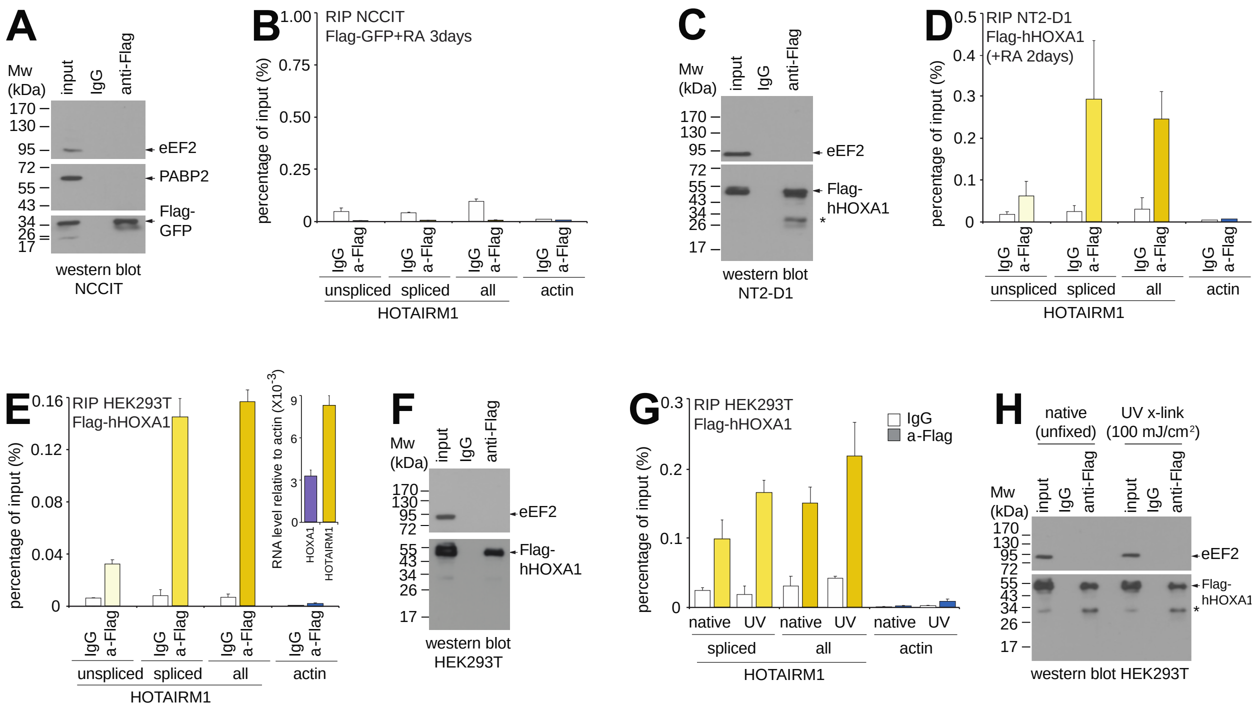


**Supplemental Figure 4.** HOTAIRM1 specifically associates with the HOXA1 transcription factor, irrespective of cell type, signaling pathway context, or experimental conditions. (*A*, *B*) HOTAIRM1 is not immunoprecipitated by transfected Flag-GFP in RA-induced NCCIT cells. Western blot (*A*) shows specific immunoprecipitation of the tagged GFP protein. (*B*) Neither HOTAIRM1 nor actin was detected by RT-qPCR in RIP samples using primer sequences listed in Supplemental Table 1. All RIP measurements are shown as a percentage of input and are from at least 3 technical replicates, where error bars represent standard deviations. (*C*, *D*) HOXA1 also binds HOTAIRM1 in RA-induced NT2-D1 cells. Western blot (*C*) analysis of the RIP samples analyzed in (*D*) shows specific pull-down of transfected Flag-hHOXA1 in NT2-D1 cells. (*D*) RIP samples were probed by RT-qPCR as in (*B*). (*E*) HOTAIRM1 also associates with HOXA1 in HEK293T cells. RNAs immunoprecipitated with either IgG or Flag antibodies were measured as in (*B*). Inset shows levels of HOXA1 and HOTAIRM1 RNA in HEK293T cells relative to actin. (*F*) Flag-hHOXA1 transfected in HEK293T is specifically immunoprecipitated. Western blot analysis of the input (1%) and RIP (5%) samples from the RIP presented in (*E*). (*G, H*) HOTAIRM1 associates with HOXA1 regardless of if cells are first UV cross-linked. RNAs immunoprecipitated with transfected Flag-hHOXA1 were measured as in (*B*). In all western blots, inputs represent 1% of the material used in RIPs, and IgG or anti-Flag lanes correspond to 5% of the RIP samples. Asterisks denote antibody background signal.

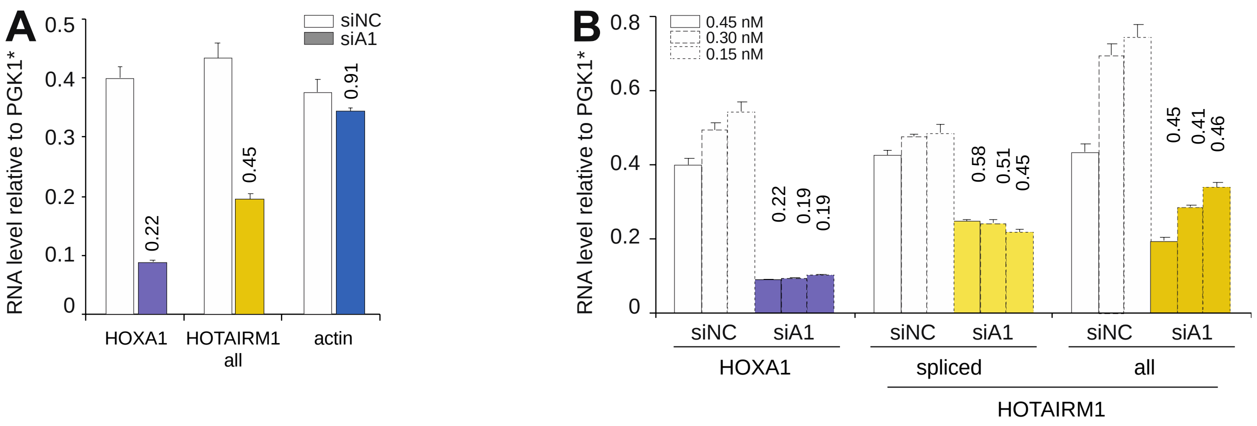


**Supplemental Figure 5.** Depleting HOXA1 blunts HOTAIRM1 induction during RA-mediated NT2-D1 differentiation. (*A*) Biological replicate of Fig. 5C confirms lower HOTAIRM1 mRNA levels when HOXA1 is RNAi-depleted during NT2-D1 differentiation (3 days post-RA treatment). HOTAIRM1, but not actin levels, were specifically affected by HOXA1 depletion. *HOXA1 values were divided by 5 and actin values divided by 100 to display on the same scale with HOTAIRM1. (*B*) Lower HOTAIRM1 levels in HOXA1 knockdown cells is not an off-target effect of RNAi depletion. HOXA1 was depleted using three different siRNA concentrations (siA1) and NT2-D1 cells were collected 3 days post-RA. mRNA levels were measured against control cell samples treated with corresponding siNC concentrations, with numbers above histogram bars representing fold differences. *HOXA1 values were divided by 5 to display on the same scale with HOTAIRM1. All RNA levels are relative to PGK1 and measured by RT-qPCR. Expression values are averages from at least 3 measurements and error bars denote standard deviations.

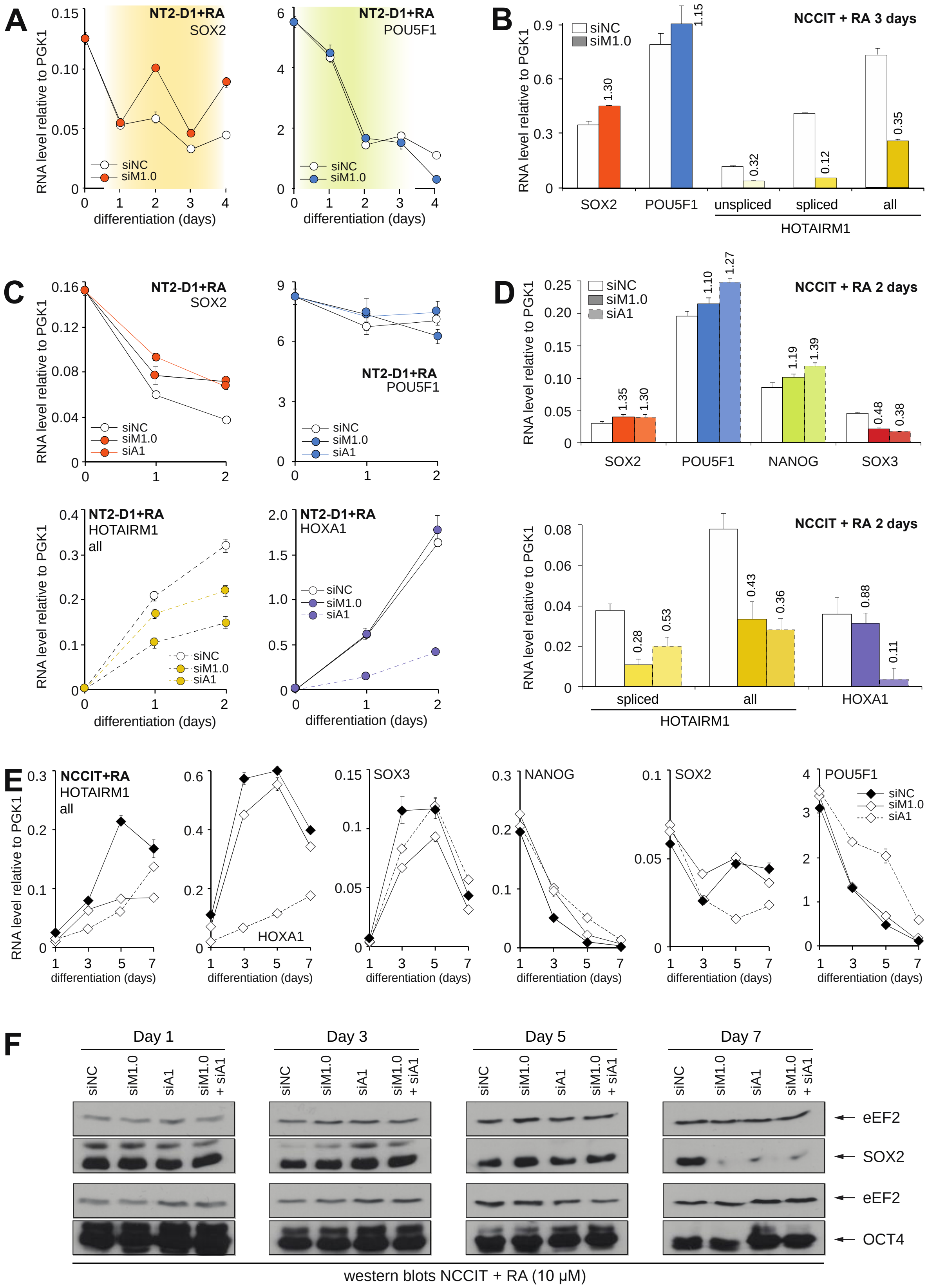
A
NT2-D1+RA
SOX2
RNA level relative to PGK1
siNC
siM1.0
differentiation (days)
NT2-D1+RA
POU5F1
differentiation (days)
B
NCCIT + RA 3 days
siNC
siM1.0
RNA level relative to PGK1
1.30
1.15
0.32
0.12
0.35
SOX2
POU5F1
unspliced
spliced
all
HOTAIRM1
C
NT2-D1+RA
SOX2
siNC
siM1.0
siA1
NT2-D1+RA
POU5F1
NT2-D1+RA
HOTAIRM1
all
NT2-D1+RA
HOXA1
differentiation (days)
D
NCCIT + RA 2 days
siNC
siM1.0
siA1
1.35
1.30
1.10
1.27
1.19
1.39
0.48
0.38
SOX2
POU5F1
NANOG
SOX3
NCCIT + RA 2 days
0.28
0.53
0.43
0.36
0.88
0.11
spliced
all
HOXA1
HOTAIRM1
E
NCCIT+RA
HOTAIRM1
all
HOXA1
SOX3
NANOG
SOX2
POU5F1
siNC
siM1.0
siA1
differentiation (days)
F
Day 1
Day 3
Day 5
Day 7
siNC
siM1.0
siA1
siM1.0 + siA1
eEF2
SOX2
eEF2
OCT4
western blots NCCIT + RA (10 µM)

**Supplemental Figure 6.** *SOX2* repression is more sensitive to HOTAIRM1 depletion than *POU5F1* is during early neuronal differentiation. (*A*) Biological replicate time-course of Fig. 7B,E over an extended time frame, confirming that depleting HOTAIRM1 during NT2-D1 differentiation delays *SOX2* repression while not considerably affecting POU5F1 levels. HOTAIRM1 knockdown levels and HOXA1 expression are found in Supplemental Fig. 3D. (*B*) Same as in (*A*) except that differentiation was in NCCIT and cells were collected only after 3 days. HOTAIRM1 knockdown levels and HOXA1 expression are found in Fig. 6A. (*C*) Additional biological replicates of RNAi depletion time-courses of HOTAIRM1 or HOXA1 during NT2-D1 differentiation by RA. (*D*) Same as (*C*) except that differentiation was in NCCIT and cells were collected after 2 days. (*E*) Biological replicate time-course of control, HOTAIRM1 and/or HOXA1 siRNA knockdowns in NCCIT cells. (*F*) Western blot analysis of SOX2 and OCT4 protein levels corresponding to samples characterized by RT-qPCR in (*E*). All RNA levels are relative to PGK1 and measured by RT-qPCR. Expression values are averages from at least 3 measurements and error bars are standard deviations.

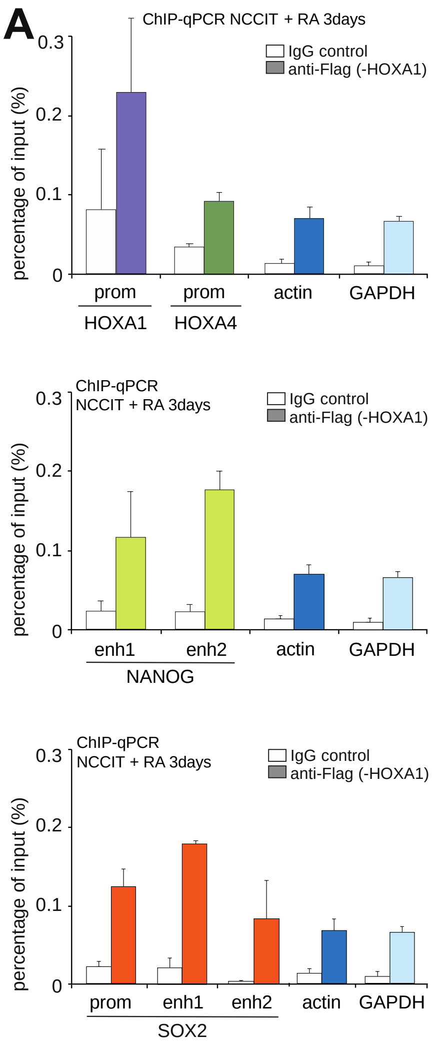


**Supplemental Figure 7.** HOXA1 binds at the *HOXA*, *NANOG*, and *SOX2* loci in human cells. (*A*) Biological replicate of the ChIP-qPCR analysis of transfected Flag-hHOXA1 in NCCIT treated 3 days with RA (10 µM). Regions in the actin and GAPDH genes were used as negative controls and shown in each panel for direct comparison. The genomic positions of PCR amplicons are found in Fig. 8D, and primer sequences used are in Supplemental Table 1. Error bars correspond to standard deviations of at least 3 technical replicates.
